# Apurinic/apyrimidinic endodeoxyribonuclease 1 contributes to the repair of damaged intercalated-motif of telomeric sequences

**DOI:** 10.64898/2025.12.17.694817

**Authors:** Alessia Bellina, Matilde Clarissa Malfatti, Tobias Obermann, Kayla Mae Grooms, Andreas Gjøsæther, Zahraa Othman, Gilmar Salgado, Daniela Marasco, Antonella Virgilio, Veronica Esposito, Giulia Antoniali, Catia Mio, Matteo Pivetta, Magnar Bjørås, Barbara van Loon, Gianluca Tell

**Author notes:** Correpondence may also be addressed to Matilde Clarissa Malfatti.

## Abstract

Apurinic/apyrimidinic endodeoxyribonuclease 1 (APE1) is a key enzyme in the Base Excision Repair pathway, responsible for processing abasic (AP-) sites. Recent studies revealed that APE1 participates in repairing DNA secondary structures as G-quadruplexes (G4). Telomeres, stabilized by shelterin proteins, are rich in G4, where APE1 binds and repairs AP-sites to maintain their integrity. The complementary cytosine-rich strand forms another structure, the i-motif (iM), essential for telomere maintenance, though its repair mechanism remains unclear.

Herein we investigate APE1 binding and processing capabilities toward native and damaged telomeric iM, bearing AP-sites in different positions. Using biochemical and biophysical assays, we found that APE1 binds the telomeric iM-sequence and that its cleavage efficiency depends on AP-site position within iM. Proximity Ligation Assay analysis, in HeLa and U2OS cells, highlighted a novel interaction between APE1 and PCBP1, a well-known iM-folding modulator. PCBP1 binds iM with higher affinity than APE1 and inhibits its cleavage activity on damaged iM. Immunofluorescence and Telomere Restriction Fragment analyses showed that depletion of APE1 or PCBP1 impairs their interaction with the shelterin components, affecting telomere length. These results connect APE1 canonical DNA repair activity with the maintenance of non-canonical DNA secondary structures in telomeres, through its interaction with PCBP1.

**GRAPHICAL ABSTRACT:** 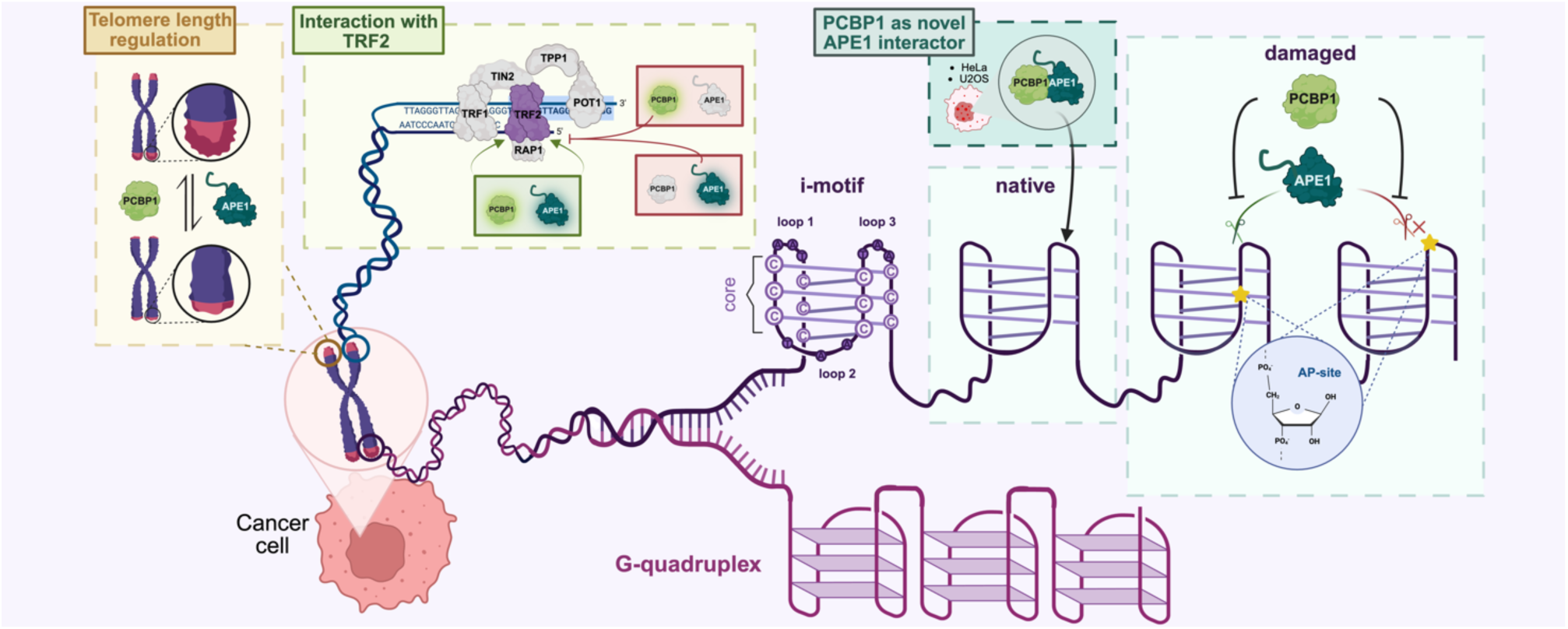

## INTRODUCTION

Intercalated motif (iM) DNA structures consist of two parallel-stranded duplexes intercalated in antiparallel orientation and held together by hemi-protonated cytosine-cytosine^+^(C:C^+^) base pairs (1). The iM core is always composed of a scaffold of C:C^+^ base pairs, which varies in length between different iM, and by the loop regions connecting the core, which are specific for each sequence, differing in length and nucleotide composition (2). It is well established that the formation of iM is dependent on acidic pH *in vitro* (1), while *in vivo* it might occurr also at physiological pH (3, 4), specifically under conditions of molecular crowding (5, 6), negative superhelicity induced by transcription (7) and in the presence of silver and copper cations (8, 9). Through the Quadparser algorithm (3), sequences having four tracts of five cytosines, separated by 1 to 19 nucleotides and able to form iM, were searched in the human genome (3), and surprisingly 5’125 potential iM sequences were identified (3), with a 12.4% of them prevalently located in gene promoters. Through a bioinformatic analysis with the R package Biostrings, it has been demonstrated the presence of 769 dCn sequences, with *n* between 15 and 81 nucleotides, mainly at promoter levels, introns and 5’ and 3’ UTR (4). Moreover, by using a specific synthetic antibody fragment (iMab), which recognizes iM with high selectivity and affinity in cells, it has been discovered that the number of iM foci dynamically changes during cell cycle progression (10). Specifically, the highest number of foci formation occurs during the G1/S phase boundary with a consequent decrease in S-phase, suggesting that iM are resolved before DNA replication with a role in regulating transcriptional activity (10). The most relevant demonstration of an iM-regulatory role on DNA transcription concerns *BCL2* (11) and *HRAS* oncogenes (12). Regarding *BCL2*, its promoter folds into a harpin or an iM, which are in a dynamic equilibrium. The ribonucleoprotein hnRNP-L binds to its iM, causing its unfolding to a single strand thus leading to the activation of the *BCL2* gene transcription (11). Similarly, the ribonucleoprotein hnRNP A1 is able to unfold the iM present in the promoter of *HRAS*, leading to the activation of the *HRAS* gene transcription (12). Many studies demonstrated the presence of iMs also within telomeres (10, 13, 14), which are the protective ends of the chromosomes formed by 5-15 kb long tracts of double-stranded guanine and cytosine repeats, with a 50-300 nucleotides protrusion of single-stranded guanine repeats at the 3’end, called G-tail or G-overhang (15, 16). Under physiological conditions, telomeres shorten every cell division until reaching a critical length that triggers intracellular senescence signals (16, 17). This highly repetitive and GC-rich nature of telomeres is strictly tied with their capability of forming high-order DNA secondary structures, such as G-quadruplex (G4) and iM (18). In detail, in U2OS cell line, a co-localization signal between the shelterin-complex protein Telomeric repeat-binding factor 2 (TRF2) and iM foci was observed (10), confirming the presence of iM structures at the telomeric level, with potential relevant biological roles. After a treatment with an iM stabilizing ligand (*i.e.* Single-walled carbon nanotubes - SWNT), the activity of telomerase was hindered, preventing the extension of telomeric DNA and leading to defective telomere maintenance (13). In addition, this treatment caused telomere uncapping, characterized by high levels of anaphase bridges and micronuclei, the release of telomere-binding proteins as TRF2 and Protection of telomeres protein 1 (POT1), which induces telomere malfunction and DNA damage signaling, with a consequent phenotype characterized by cell cycle arrest, apoptosis and senescence in HeLa cells (14). *In vitro* experiments demonstrated that both unstructured telomeric C-strands and telomeric iM inhibit the processivity of telomerase extension of parallel G4 and linear telomeric DNA (19) and recently, it has been proposed that iM structural dynamics can be modulated by a specific protein family, called poly(C)-binding proteins (PCBPs) (11, 12, 20–22). In particular, one member of this family, PCBP1 (also called hnRNP E1) was already reported to be involved in the competitive formation of iM and G4 structures in NMuMG and A549 cell lines and in the maintenance of genomic integrity. Specifically, PCBP1 co-localized with iM and its knockdown led to a decreased accumulation of iM foci in the nucleus and the induction of DNA damage signaling, which is very high upon genotoxic treatments (23). Furthermore, cell treatment with SWNT influenced PCBP1 localization, delocalizing it from telomeres as TRF2 and POT1 (14).

At present, there is no evidence whether lesions occurring on iM may affect their structural and functional properties and, above all, which proteins are responsible for the repair of damaged iM DNA structures. Apurinic/apyrimidinic (AP-) sites are among the most frequent lesions in cells, as they happen almost 10’000 times *per* human cell *per* day (24). They can be generated spontaneously or enzymatically by the action of DNA glycosylases on damaged bases such as oxidized bases (24). AP-sites are recognized and processed by the apurinic/apyrimidinic endodeoxyribonuclease 1 (APE1), the principal enzyme of the Base Excision Repair (BER) pathway (25). Several studies, performed by our laboratory and others, showed that APE1 holds a highly substrate- and structure-dependent endonuclease activity (26, 27), particularly in the context of non-canonical secondary structures. Indeed, APE1 activity is lower on damaged telomeric G4 structures, compared to the canonical double-stranded substrates, and strongly depends on: i) the position of the damaged base, ii) the N-terminal region of the protein and iii) the ionic strength of the reaction (28–30). Differently from G4 structures, which have been widely studied in the last years (28–31), the role of APE1 on their iM counterparts has not been investigated, yet. Recently, Dvorakova and colleagues studied how natural base lesions (i.e. AP-sites, uracil, 8-oxo-adenine) impact telomeric iM formation, revealing that modifications in the loops may cause minor alterations in the iM formation and stability, while modifications in the core have a more extensive effect, with different consequences depending on their position and abundance (32). On this basis, this study was aimed at characterizing the effects of AP-sites on the folding of telomeric iM structures, the processing activity of APE1 on damaged structures and the effect of PCBP1 protein on APE1 activities. Using complementary biochemical, biophysical and cellular investigations, we showed that: i) APE1 is able to stably bind the telomeric iM structure; ii) AP-containing telomeric sequences can fold into iM structures and AP-sites affect the iM-folding and stability according to their position in the structure; iii) APE1 cleavage activity on AP-sites strongly depends on the AP-position and on APE1 N-terminal region; iv) APE1 and PCBP1 are found in close proximity in HeLa and U2OS cellular lines; v) PCBP1 inhibits APE1 cleavage on telomeric iM sequence; vi) the depletion of APE1 and PCBP1 dysregulates telomere length with opposite trends in a Alternative Lengthening of Telomeres (ALT)-U2OS model; vii) APE1 and PCBP1 can mutually and dependently interact with the shelterin protein TRF2; viii) APE1 depletion correlates with a diminishment of iMab foci in U2OS cells; ix) APE1 and PCBP1 depletion leads to DNA Damage Response (DDR) activation in U2OS cellular model, with a possible radical alteration of DNA damage signaling when they are both depleted. Our study, besides deeply characterizing a still unexplored field in DNA repair mechanisms of BER enzymes, opens novel perspectives of translational applications in biology and medicine, which far extend BER function in telomere biology.

## MATERIAL AND METHODS

### Protein expression and FPLC purification

*E. coli* BL21 (DE3) bacteria (C2530H, New England Biolabs) were transformed with 100 ng of plasmids pGEX-3X APE1^WT^, pGEX-3X APE1^NΔ33^, pTAC-MAT-APE1^WT^-GFP, pTAC-MAT-APE1^NΔ33^ -GFP or pET166-E1 for the expression of GST-tagged APE1^WT^, APE1^NΔ33^, His- and GFP-tagged APE1^WT^, APE1^NΔ33^ and His-tagged PCBP1, respectively. The bacterial culture was harvested at 37°C at 250 rpm. After reaching an OD600 = 0.6, the protein expression was induced with 1 mM IPTG (11411446001, Roche) and bacteria were left to grow for 4 hours at 37°C. Pellets were then lysed in the presence of a protease inhibitors cocktail (P8465, Merck) and lysozyme through six cycles of sonication of 30 sec. The sample was then centrifuged at 23’000 x g for 20 minutes at 4°C and the supernatant was conserved and filtered with a 0.45 µm filter for the following FPLC purification (GE Healthcare). Purification of the GST-tagged recombinant proteins was carried out using GSTrap columns (GE17-5281-01, GE Healthcare) following the manufacturers instructions. Protein elution was obtained by 10 mM GSH (G4251, Merck). The GST-tag was also removed through incubation with Factor Xa (F9302, Merck) following the manufacturers instructions and the recombinant protein was separated from the tag by using a HiTrap Benzamidine HP column (GE17-51243-01, GE Healthcare). The purification of the His-tagged recombinant proteins was carried out using an HisTrap HP (IMAC(Ni^2+^)) column (GE17-5247-01, GE Healthcare) and were eluted by setting an imidazole gradient, with a final concentration of 800 mM. All protein fractions were dialyzed and stored at -80°C in 25 mM Tris-HCl pH 7.5, 100 mM NaCl, 10% glycerol.

### Oligonucleotides and annealing

All the oligonucleotides (ODNs) used in this study are reported in Table 1. The ODNs were synthesized by Metabion and Ella Biotech, purified by HPLC and checked in Mass Spectrometry. Where indicated, a tetrahydrofuran base, mimicking an abasic site (AP), substitutes the canonical nucleotide. Where specified, the ODN holds an IRDye-800 or a quencher BBQ650 at the 5’. The ODNs were resuspended in DNase- and RNase-free water, at a final concentration of 100 µM. To allow iM formation, 100 pmol of each ODN was prepared in a final volume of 40 μL with 50 mM Tris-Acetate, pH 5.5, and 50 mM KCl and allowed to sit o/n at RT. For the experiments with double-stranded ODNs, 100 pmol of each ODN were annealed with 150 pmol of its complementary DNA oligonucleotide in 10 mM Tris-HCl pH 7.4 and 10 mM MgCl_2_, heated at 95°C and cooled down o/n in the dark. For the Lumicks C-trap approach, DNA iM substrate, called 3x-C-NAT, was purchased from Ella Biotech and its sequence is reported in Table 1. The substrate was ligated inside long handles by using the hairpin labeling and tethering kit (Lumicks, 00016/17). Long handles are derivatized with three biotins and three digoxigenins at the opposite ends, to allow the tethering by using streptavidin and anti-digoxigenins beads.

**Table 1:**
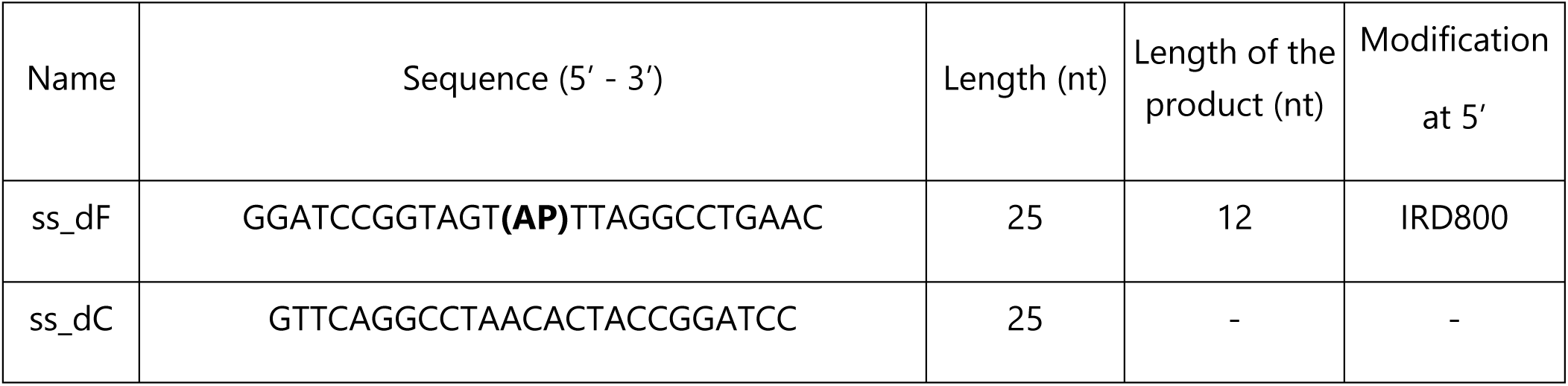
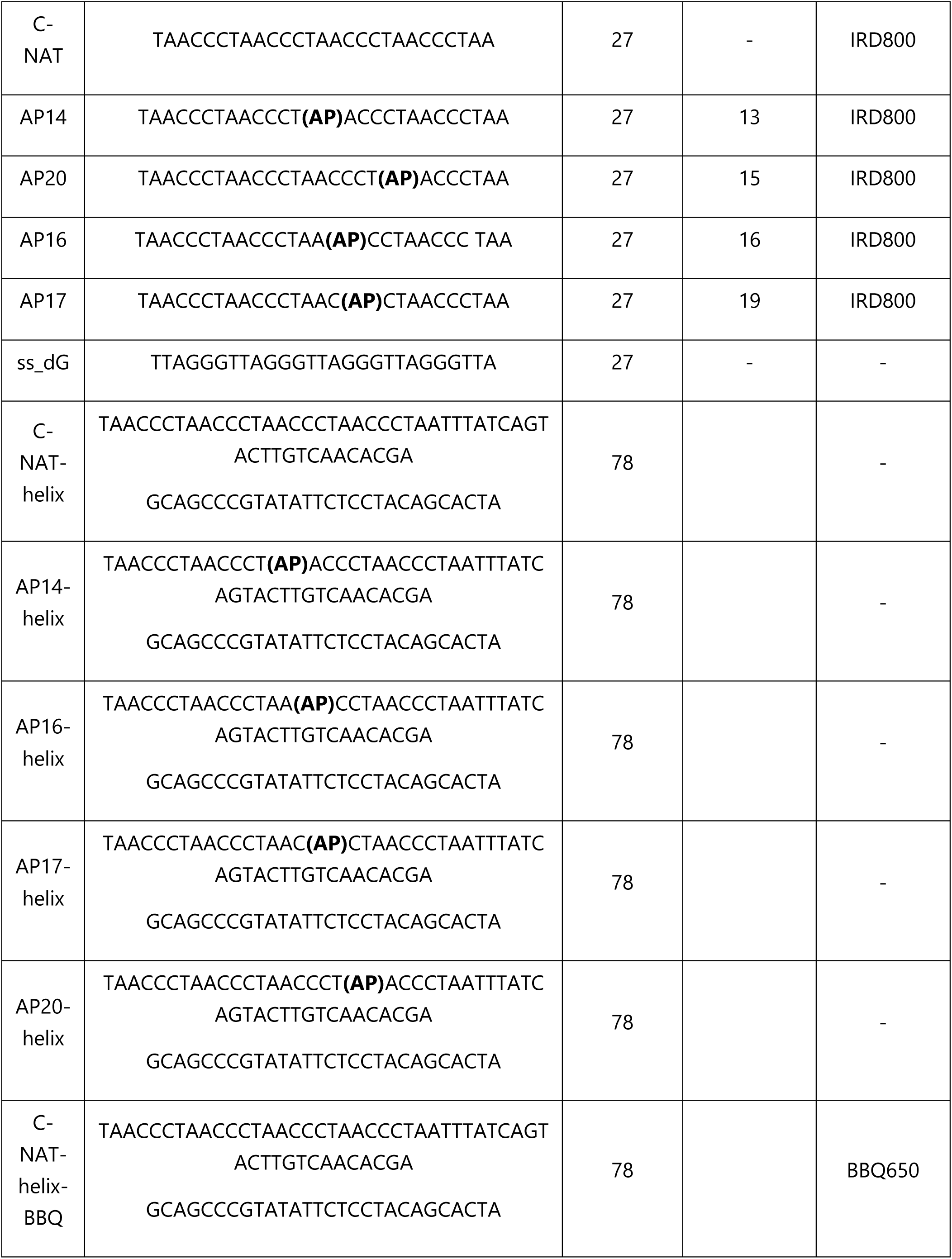
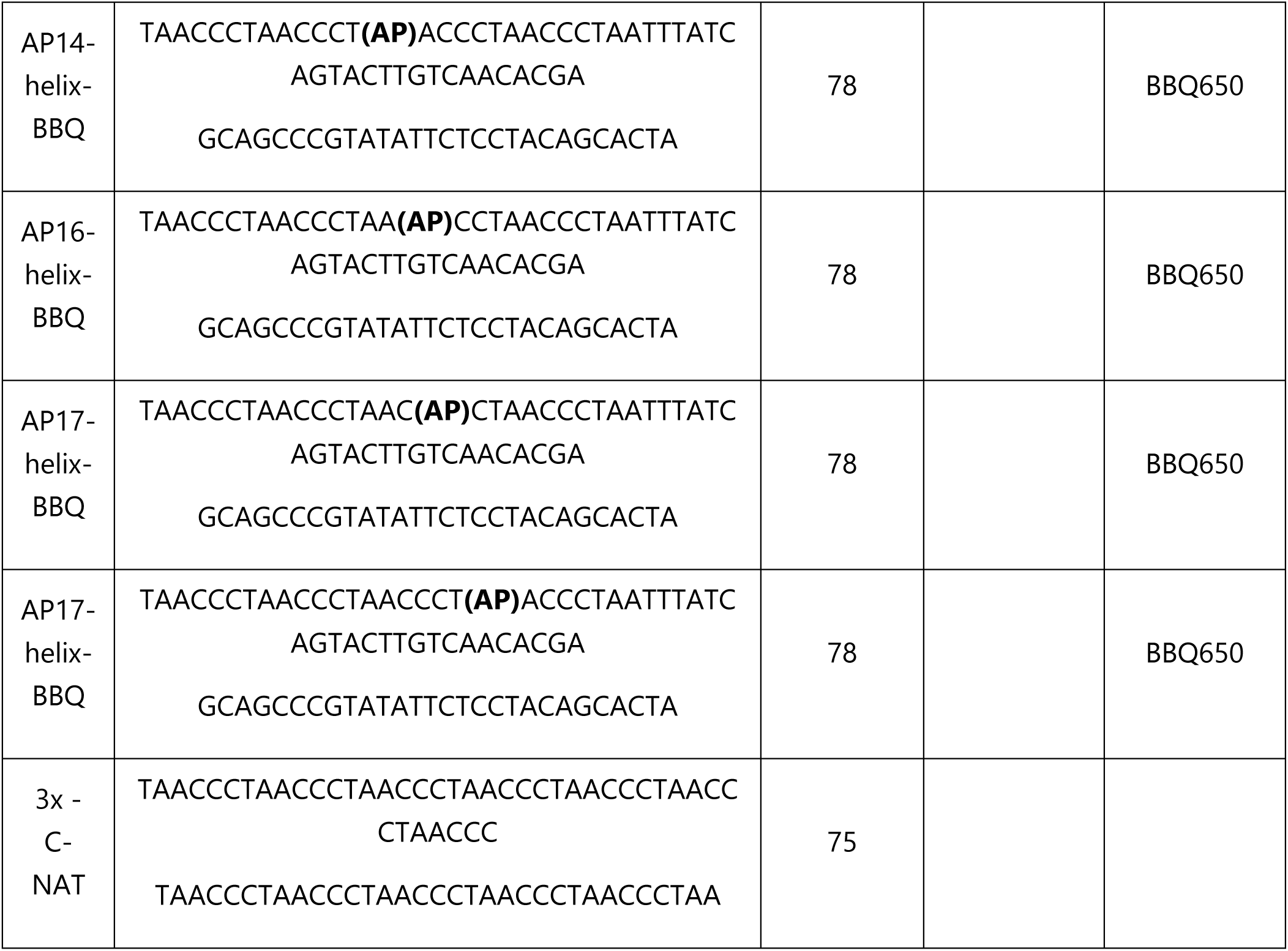
Sequences of the ODNs used in this study. The name of the ODNs is reported in the first column. The sequence of the 5’ - 3’ single-stranded (ss) ODNs is reported in the second column. The modified bases are highlighted in bold type. The whole length and the length of the product are reported in the third and fourth columns. In the fifth column, the modification at the 5’ of each ODN is reported.

### Circular dichroism (CD) spectroscopy

CD samples of ODNs reported in Table 1 were prepared at a ODN concentration of 6 µM by using a buffer with 50 mM Tris-Acetate, 50 mM KCl, pH 5.5 and submitted to the annealing procedure by heating at 70°C and slowly cooling at room temperature. CD spectra of all iM structures and CD melting curves were registered on a Jasco 715 CD spectrophotometer (Jasco, Tokyo, Japan). For the CD spectra, the wavelength was varied from 360 to 220 nm at 100 nm min^-1^ scan rate, and the spectra were recorded with a response of 4 s, at 2.0 nm bandwidth and normalized by subtraction of the background scan with buffer. The temperature was kept constant at 10°C with a thermoelectrically-controlled cell holder (PTC-348, Jasco). CD melting curves were registered as a function of temperature from 10°C to 80°C for iM-forming ODNs at their maximum Cotton effect wavelengths. The CD data were recorded in a 0.1 cm pathlength cuvette with a scan rate of 0.5°C/min.

### NMR spectroscopy

NMR samples were prepared in buffer containing 50 mM NaCl, 10 mM MES (pH 5.5) and 1 mM DTT, unless otherwise indicated. All measurements were acquired at 25 °C in 3 mm thin-wall NMR tubes with 5% D2O. Recombinant APE1 was uniformly 15N-labeled and dissolved in the same buffer at a concentration of 100 µM. The C-NAT oligonucleotide was dissolved in identical buffer and annealed by heating to 95 °C for 5 min followed by slow cooling to ambient temperature. For titration experiments, the C-NAT was added stepwise to the 15N-labeled UP1 solution at defined molar ratios.

NMR spectra were recorded on a Bruker 800 MHz AVANCE NEO spectrometer (IECB, Bordeaux, France) equipped with a TCI cryoprobe. One-dimensional 1H spectra were acquired using the zgesgppe pulse sequence, and two-dimensional 1H–15N correlation spectra were obtained with the trosyett3gpsi sequence. Spectra were collected for APE1 alone and after successive additions of the C-NAT. Data were processed with TopSpin 4.1 (Bruker Biospin) using standard procedures. Chemical shift perturbations were calculated as weighted combined differences in 1H and 15N shifts according to the equation:

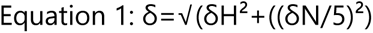

Assignments were transferred from published APE1 datasets (PDB code : 1bix), and perturbation analyses were used to identify residues involved in i-motif binding, with δH the shift in 1H dimension and δN the shift in 15N dimension. The most significate shifts were depicted on the surface of APE1 using UCSF ChimeraX.

### SwitchSENSE technology

SwitchSENSE experiments were performed on a HeliX^®^ instrument (Dynamic Biosensors GmbH) on standard switchSENSE adapter chips (ADP-48-2-0, Dynamic Biosensors GmbH) using the static measurement mode. The experimental workflow was designed in the HeliOS software (Version 2024.2.1, Dynamic Biosensors GmbH). To investigate the interaction between iM and APE1, the adapter chips were first functionalized. On the chip surface, a DNA construct called nanolever is attached. The part of the nanolever that is connected with the chip is the anchor strand and is extended by the adapter strand. The 3’ end of the adapter strand is derivatized with a fluorescent dye (Ra). This dye is used as the read out and is excited in the range of 600-630 nm and emission was recorded in the range of 650-680 nm. The complementary sequence of the adapter strand is annealed with its counterpart which towards the 5’ end is extended by our sequences of choice. Here, two new sets of ODNs (Table 1) were synthesized (Ella Biotech). One set of ODNs was modified with a quencher (BBQ650) at the 5’ end of the oligonucleotide and was exclusively used for iM-folding experiments. To account for unspecific binding of APE1 towards double-stranded DNA (dsDNA), the adapter chips have a reference spot. Here, the nanolever set-up is similar, despite that no overhang was designed. Hence, the nanolever is in dsDNA configuration and the signal recorded serves as a reference. Also, for the iM-folding experiments were the quencher was employed, the nanolever was in a dsDNA configuration, to account for pH dependent quenching.

iM-forming ODNs were hybridized with Adapter strand 1-Ra at 1:1 ratio (v/v), trough incubation at 25°C for 30 minutes at 650 rpm. For the reference, adapter strand 2, pre hybridized with the complementary sequence, was then added at a 1:1 ratio (v/v). Experiments were carried out in MES200 buffer (10 mM MES; 200 mM NaCl; 50 µM EDTA; 50 µM EGTA; 0.05 % Tween-20), at a pH of 5.5 or else specified. Two different principal approaches were used for the SwitchSENSE experiments. The first one relies on the iM recognition by a ligand, which leads to changes in the dye environment and in the fluorescence signal. For the iM folding validation, the used ligand was the iMab antibody (10). For the binding assays, APE1^WT^-GFP and APE1^NΔ33^-GFP were used as ligands. Here, the different ligands have been applied in different concentrations to obtain a dose dependent response curves. Additionally, blanks containing only the MES200 buffer were ran before and after the ligand runs. After every concentration the chip surface was regenerated using 6M guanidinium hydrochloride, subsequently the chip was functionalized with a new nanolever. Thereby we ensured that all ligand is stripped off from the chip surface. To be able to obtain the desired kinetic values, every measurement consist of an association phase and dissociation phase. During the association phase the respective ligand is injected onto the chip at a flow rate of 200 µl/min. In the dissociation phase the chip surface is washed with the MES200 buffer at a flow rate of 500 µl/min. The chip temperature is set to 25°C, to ensure ligand activity, the sample tray was cooled to 10°C.

The second approach relies on the presence of BBQ650 at the 5’ end of the oligonucleotide. Upon iM folding, the quencher is physically close to the dye of the complementary strand and induces a quenching signal. By subsequently injecting buffers with increasing pH (association phase), it can be determined if the quencher is still in close proximity to the dye. After having determined the iM formation by dye quenching, the unfolding is induced by injecting a high pH solution, consequently the quencher is led far away, which is represented by fluorescence increase. Parameters during the association and dissociation phase are as described before. Chip temperature was as well set to 25°C, and sample tray was at room temperature.

To analysis the obtained fluorescence signals were referenced against spot 2 to results in a kinetic curve representing ligand-iM interaction without unspecific binding to the double-stranded nanolever. Additionally, the blanks were used to set a stable baseline. The chosen fit model was individually chosen, based on the generated data.

In case of the quencher experiments, spot 2 served for a reference for pH-dependent quenching. Here, we were not generating kinetic data, thus no fit model was applied. iM formation was measured in % Fluorescence change.

### SDS-PAGE, Western Blot and Southwestern Blot (SWB) analysis

Recombinant proteins and cell extracts were loaded on SDS-PAGE gel for electrophoresis. For the quantification of recombinant proteins, each band was quantified and normalized with a known concentration BSA (bovine serum albumin) standard curve. The staining of the gel was performed with Coomassie Blue.

For the Western Blot analysis, upon running the gel, the proteins were transferred to a nitrocellulose membrane (Amersham^TM^ Protran®0.2 µm NC, GE10600001, Merck). Protein transferring was allowed in Towbin buffer (trizma 25 mM, glycine 23 mM, methanol 20% (v/v)) at constant 70 V, for 3 hours at 4°C. The membrane was colored with Revert 700 Staining (926-11010, Li-Cor Biosciences) and acquired with Odyssey CLx scanner/ImageStudio Software (Li-Cor Biosciences). The membrane was blocked for 1 hour with 5% BSA in PBS-Tween 0, 1% and then incubated with primary antibody for 3 hours at RT. The primary antibodies used and their dilution usage were the following: APE1, mouse, monoclonal, 1:2’000, NB 100-116, Novus; APE1, rabbit, polyclonal, 1:2’000, NB 100-101, Novus; PCBP1, mouse, 1:500, sc-137249, Santa Cruz Biotechnology; PCBP1, rabbit, polyclonal, 1:2000, NBP2-55063, Novus ; Tubulin, mouse, monoclonal, 1:2’000, T0198, Merck; p21, rabbit, polyclonal, 1:500, 2947, Cell Signaling; 𝛾H2AX, mouse, monoclonal, 1:500, 05-636, Merck; TRF2, mouse, monoclonal, sc-271710, Santa Cruz Biotechnology; HSP70, rabbit, polyclonal, 1:2’000, GTX111088, GeneTex. Then, the membrane was incubated with secondary antibodies labeled with IRDye 700 or 800 (1:10’000, 926-68071, Li-Cor Biosciences).

For the Southwestern Blot (SWB) analysis, 2 µg of each protein was loaded on the SDS-PAGE gel. After running and blotting, the membrane was incubated for 10 minutes at room temperature in a 6M guanidine hydrochloride solution to denature the proteins. The proteins were then renatured by incubating the membrane with serial guanidine hydrochloride dilutions until 0.094 M in Southwestern Blot Buffer (SWB Buffer, 10 mM HEPES, 50 mM NaCl, 10 mM MgCl2, 0.1 mM EDTA, 1 mM DTT, 50 μM ZnSO4, 0.1% Tween-20) for 10 minutes at 4°C. The membrane was then washed twice with SWB buffer and then incubated for 1 hour at room temperature with SWB blocking buffer (5% BSA in SWB buffer). At the end, the membrane was incubated with 5 pmol of the fluorescent oligonucleotide in SWB blocking buffer at 4°C with gentle rocking o/n, washed with SWB buffer and analyzed.

All images were acquired by Odissey CLx scanner (Li-Cor Biosciences) and analyzed by ImageStudio Software (Li-Cor Biosciences).

### AP-site incision assay

To measure endonuclease activity of recombinant proteins, we performed AP-site incision assay as described in (33, 34). Briefly, the oligonucleotide substrates (25 nM) were incubated at 37°C with different amounts of proteins and for different timing points, as indicated in the legend of each figure. Reactions were carried out in a buffer containing 20 mM TrisHCl pH 7.4, 50 mM KCl, 1 mM MgCl2, 0, 1% BSA(v/v), 0, 1% Tween-20 (v/v). At the end, reactions were blocked with a stop solution (99.5% (v/v) formamide, 10× Orange Loading Dye (927-10100, Li-Cor Biosciences)) and heated at 95°C for 5 min. Samples were loaded onto a 7 M urea denaturing 20% polyacrylamide gel in TBE buffer (45 mM Tris, 45 mM Boric Acid, 1 mM EDTA, pH 8.0) and run at 4°C at 300 V for 30 minutes. The gel was directly scanned by Odyssey CLx scanner (Li-Cor Biosciences) and analyzed by ImageStudio Software (Li-Cor Biosciences).

### Cell culture, CRISPR/Cas9, siRNA transfection and nuclear cell extract preparation

For the experimental procedures performed in mammalian cells, different human tumoral adherent cell lines were used including HeLa cells, from cervical tumor, U2OS cells, from bone osteosarcoma and A549 clone 31, from lung adenocarcinoma. HeLa and U2OS were grown in DMEM High Glucose medium (ECB7501L, EuroClone), complemented with 10% (v/v) of fetal bovine serum (FBS), penicillin (100 U/mL), streptomycin (100 mg/mL) and l-glutamine (2 mM) (EuroClone). A549 clone 31 cells were grown in RPMI 1640 medium (ECM9106L, EuroClone), complemented with 25 mM Hepes, 10% (v/v) FBS, penicillin (100 U/mL), streptomycin (100 mg/mL) and l-glutamine (2 mM) (EuroClone). All cells were grown in an incubator at a constant and monitored temperature of 37°C and partial pressure of CO_2_ of 5%.

The A549 clone 31 cell line was generated using the strategy previously described in (35) and deeply characterized in the Supplementary section of this article. Briefly, APE1^WT^-mEGFP-expressing cells were generated by transiently transfecting A549 cell line with Lipofectamine 3000 (L3000001, Life Technologies) through puC19 repair template for the tagging of APE1 C-term at endogenous loci together with two plasmids encoding for two single guide RNAs (sgRNAs), following the manufacturer’s instructions. Cells were expanded for 7 days before FACS sorting with Multi application cell sorter MA900 (Sony Biotechnology Inc.) based on the mEGFP signal. All cells mEGFP^+^ were then clonally expanded and characterization of single-cell clones was performed by immunoblotting and Sanger sequencing.

For the HeLa and U2OS silencing, 300’000 cells were plated in a six-multiwell. The day after, cells were transfected with 100 pmol of scramble (5’ - CCA UGA GGU CAG CAU GGU CUG UU - 3’, siSCR), as negative control, 100 pmol of ON-TARGETplus Human PCBP1 SMART pool (siPCBP1), 100 pmol of siAPE1 (5’ – UACUCCAGUCGUACCAGACCU - 3’) (T-2005-01, Dharmacon), with 2.5 μL of Dharmafect (Dharmacon) in 200 μL of OPTIMEM (3185070, ThermoFisher) for 6 hours at 37°C. Next, cells were added with 1 mL of fresh DMEM, without removing the complexes. After 24 hours, cells were washed once with PBS 1X and left growth in fresh medium.

Nuclear cellular extracts (NCE) with endogenously tagged APE1-mEGFP were obtained from A549 clone 31 cell line, starting from 4 million cells, and by using Nuclear extraction kit (ab113474, Abcam) by including three cycles of 10 sec sonication with Bioruptor Pico at 4°C. The resulting nuclear extracts were aliquoted into single-use tubes and flash frozen in liquid nitrogen, for then long-term storage at -80°C. NCE total protein concentration was determined by using Bradford assay (5000201, Bio-Rad), while NCE nucleic acid concentration was determined using Quant-iT™ PicoGreen™ dsDNA Assay Kit (P7589, Invitrogen). The DNA concentration in the NCE was in range of 40 ng/µl.

### Single-molecule analysis of DNA-binding proteins from nuclear extracts

SMADNE (single-molecule analysis of DNA-binding proteins from nuclear extracts) analysis was carried out on Dymo C-trap^®^ instrument (LUMICKS), by applying general principles described in (36). C-trap^®^ technology unites a microfluidic system, a dual-tap optical system and a three-color confocal fluorescence microscope. All five channels of the microfluidic flow cell were pressurized to 0.10 bar. Prior to experiments, the channel used for NCE was passivated with casein. Reagents were diluted in recording buffer (25 mM MES pH 5.5, 150 mM NaCl, 0.1 mg/mL BSA, 1 mM DTT, 1 mM Trolox). In ascending order, channels were loaded with 1.18 μm streptavidin-coated beads, 0.81 μm anti-digoxigenin beads, ligated DNA iM substrate, recording buffer, and NCE (1:45). A single DNA tether was formed between optically trapped beads by adjusting the distance between them. The tether was washed in recording buffer, flow was stopped, and force-distance curves were compared to an extensible worm-like chain model to confirm single DNA tether formation and iM structure unfolding (∼22 pN). During recordings, DNA tension was maintained between 10-15 pN, and continuous confocal line scanning was initiated along the tether as it moved into the NCE channel. The EGFP fluorescence was excited by a 488 nm laser at 5% power, detected through a 525/45 nm emission filter, and captured using a 1.2 NA, 60X water immersion objective and single-photon avalanche photodiodes. Kymographs were generated with 20 nm pixel size, 0.2 ms pixel dwell time, and 0.062 ms line time.

Kymographs were analysed using a custom Python script based on Pylake (v1.5.3, LUMICKS). Binding events were tracked using a greedy algorithm (37), by manually selecting regions encompassing individual events lasting at least 240 ms. Tracks separated by <1 s at the same position were manually connected to account for eGFP blinking. Events within 0.1 μm of the DNA midpoint (site of iM) were classified as specific. Specific gap times were defined as intervals between consecutive specific binding events. Binding and gap times were analysed by cumulative residence time distribution (CRTD) and cumulative gap time distribution (CGTD), respectively, using an exponential one-phase decay fit to obtain rate constants.

### Immunofluorescence and Proximity Ligation Assay (PLA)

For immunofluorescence analysis, 80’000 cells were plated on glass slides placed inside a 24-multiwell. After indicated treatment, cells were fixed with 4% paraformaldehyde (PFA) for 20 minutes at RT. For the detection of APE1 and PCBP1, cells were permeabilized using Triton X-100 0.25% in PBS 1X for 5 minutes at RT and blocked with FBS 10% in Wash Buffer A (WBA, 0.15 M NaCl, 0.01 M Tris, 0.05% m/v Tween 20, pH 7.5), for 30 minutes at RT. The slides were then incubated with the primary antibody (PCBP1, 1:50, sc-137249, Santa Cruz Biotechnology; APE1, 1:100, NB 100-101, Novus) diluted in FBS 10% in WBA at 37°C for 2 hours. Next, cells were incubated with the secondary antibody (Alexa Fluor 488, 111-545-003, Jackson ImmunoResearch; Alexa Fluor 633, A211050, A-21071, Invitrogen; Alexa Fluor 555, ab150114, Abcam), diluted in FBS 10% in WBA for 1 hour at RT. For the detection of TRF2, cells were permeabilized with Permeabilization solution (20 mM TrisHCl pH 8.0, 50 mM NaCl, 3 mM MgCl2, 300 mM sucrose, 0.5% Triton X-100) for 15 minutes at RT and blocked with BSA 1% in PBS for 1 hour at RT. The slides were then incubated with primary antibody (TRF2, 1:100, sc-271710, Santa Cruz Biotechnology) for 2 hours at RT, diluted in BSA 1% in PBS. Finally, cells were incubated with the secondary antibody (Alexa Fluor 488, 111-545-003, Jackson ImmunoResearch; Alexa Fluor 633, A211050, A-21071, Invitrogen; Alexa Fluor 555, ab150114, Abcam), diluted in BSA 1% in PBS for 30 minutes at RT. For the detection of iM in cells, we adapted the protocol published in (10). Birefly, cells were pre-fixed by adding directly to the media an equivolume of PFA 2% diluted in PBS 1X, incubated for two minutes at RT. Cells were washed with ice-cold PBS and fixed with PFA 2% for 30 minutes at 4°C. Cells were then washed with ice cold PBS and permeabilized using Triton X-100 0.1% in PBS 1X for 30 minutes at 4°C. Cells were blocked with iMab blocking buffer (BSA 2%, milk 1% in PBS 1X) o/n at 4°C. Slides were incubated with the primary antibody (iMab, 1:100, Ab01462-23.0, Absolute Antibody) o/n at 4°C. Finally, secondary antibody staining was performed (Alexa Fluor 488, 111-545-003, Jackson ImmunoResearch; Alexa Fluor 633, A211050, A-21071, Invitrogen; Alexa Fluor 555, ab150114, Abcam), diluted in PBS 1X supplemented wih 2% gelatin, 0.5% BSA and 0.1% Tween-2 for 45 minutes at RT. After each antibody incubation, cells were washed three times with ice-cold PBS 1X, complemented with 0.1% Tween-20. For the proximity ligation assay (PLA), the kit Duolink Proximity Ligation Assay (DUO92007, Merck) was used, following the manufacturer’s instructions. Briefly, following the incubation with primary antibodies, cells were incubated with PLA probes for 1h at 37°C (PLA probe anti-mouse minus, DUO92004, PLA probe anti-rabbit minus, DUO92002, Merck). Afterwards, slides were incubated for 30 minutes at 37°C with the Ligation Solution, which was followed by the incubation with the Amplification solution for 100 minutes at 37°C. For each protocol, nuclei were stained and sildes mounted with Mounting medium containing DAPI (Fluoroshield with DAPI – F6057, Merck). Images were acquired with a confocal microscope (Leica TCS SP8, Leica Microsystems).

### Telomere Restriction Fragment (TRF) Assay

Telomere length analysis was performed by using TeloTAGGG Telomere Length Assay Kit (12209136001, Merck) which exploits the telomere restriction fragment (TRF) principle. Briefly, genomic DNA was isolated from U2OS cells using the QiAmp DNA Mini kit (51304, Qiagen) and sonicated on Bioruptor Plus (Diagenode) at maximum power for four times for 30 seconds, with breaks of 30 seconds. Sonicated DNA was then digested with both Hinf I and Rsa I endonuclease enzymes for 2 hours at 37°C. After digestion, samples were loaded on a 0.8% agarose gel and run for 4 hours. The gel was incubated with an 0.25 M HCl solution for 10 minutes, followed by two consequent incubation with an alkalaline denaturation solution (0.5 M NaOH, 1.5 M NaCl). Afterwards, the gel was incubated twice with a neutralization solution (0.5 M TrisHCl, 3 M NaCl, pH 7.5) for ten minutes, blotted o/n by capillary transfer on a nylon membrane (HybondN+ Boehringer Mannheim), and UV-crosslinked (120 mJ). Lastly, DNA on the membrane was detected by hybridization with a digoxigenin-labeled telomeric probe, according to the manufacturer’s instructions. The membrane was scanned using a Molecular Imager Chemidoc XRS scanner (BioRad) with ImageLabTM software (BioRad). The profiles and the length of fragments were quantified using the WALTER web tool (38).

### Statistical analysis

The results are presented as means ± SD, and data analysis was performed with the Prism GraphPad

7.0 software. For comparison between two groups, Student t-test was used. For comparisons between multiple groups, ordinary one-way ANOVA was used. In all tests, a P-value < 0.05 is symbolized by a single asterisk (*), while a P-value < 0.01 is symbolized by two asterisks (**), < 0.001 by three asterisks (***), and < 0.0001 by four asterisks (****).

## RESULTS

### Biophysical characterization of telomeric iM

In order to perform binding studies of APE1 towards telomeric iM structure, firstly we checked the ability of a synthetic 27-mer oligonucleotide to fold into an iM structure, through different biophysical techniques and in different pHs. This sequence, named C-NAT, holds the nucleotide composition of the cytosine-rich strand of telomeres, (Fig. 1A and Table 1). Firstly, CD analysis at pH 5.5 revealed that C-NAT exhibits a positive peak at 288 nm and a negative one at 255 nm (Fig. 1B), typical of iM structures (39, 40), with a melting temperature (Tm) of 46°C (Fig. 1C). The folding at pH 5.5 was confirmed by 1D ^1^H NMR spectroscopy (Fig. 1D), in which only three characteristic and unique imino sharp-resonance peaks (15–16 ppm) can be observed, in a pattern corresponding to two symmetry-related C:C^+^ pairs, indicating the presence of intercalated, hemiprotonated cytosine: cytosine base pairs in a single stable conformation. Then we used the SwitchSENSE technology, by means of two different approaches, to further validate the iM folding (Supplementary Fig. S1A and S1C). Briefly, the first approach relies on the iM recognition by the iMab antibody (Supplementary Fig. S1A), able to specifically recognize iM structures both *in vitro* and in cells (10). Keeping the C-NAT concentration constant and increasing iMab concentrations, we obtained dose-response profiles both at pH 5.5 and 6.5. Whereas iMab could not dissociate from the iM at pH 5.5, resulting in a non-measurable dissociation constant (KD) (Supplementary Fig. S1B), at pH 6.5 the fitting of the kinetic curves in a biphasic model allowed to determine the first (KD1|1) and the second dissociation constant (KD2|2) of the iMab towards the iM structure, which resulted of 9.01 nM for the KD1|1 and of 0.70 nM for the KD2|2 (Fig. 1E). For the second approach, the C-NAT was modified to hold the BBQ650 (Supplementary Fig. S1C). By increasing the pH of the buffer from 5.5 to 6.2, the signal of the Ra-dye attached to the adaptor sequence was quenched far less due to the unfolding of the iM structure which leads the BBQ increasingly further from the dye (Supplementary Fig. S1D). The change in fluorescence signal (%) decreased in a linear way upon 0.1 pH changes, confirming the importance of the acidic pH for the structure folding *in vitro* (Fig. 1F) and demonstrating that the complete unfolding of the C-NAT is reached at pH 6. All these experiments clearly confirmed that C-NAT is able to fold into an iM structure.

**Figure 1:**
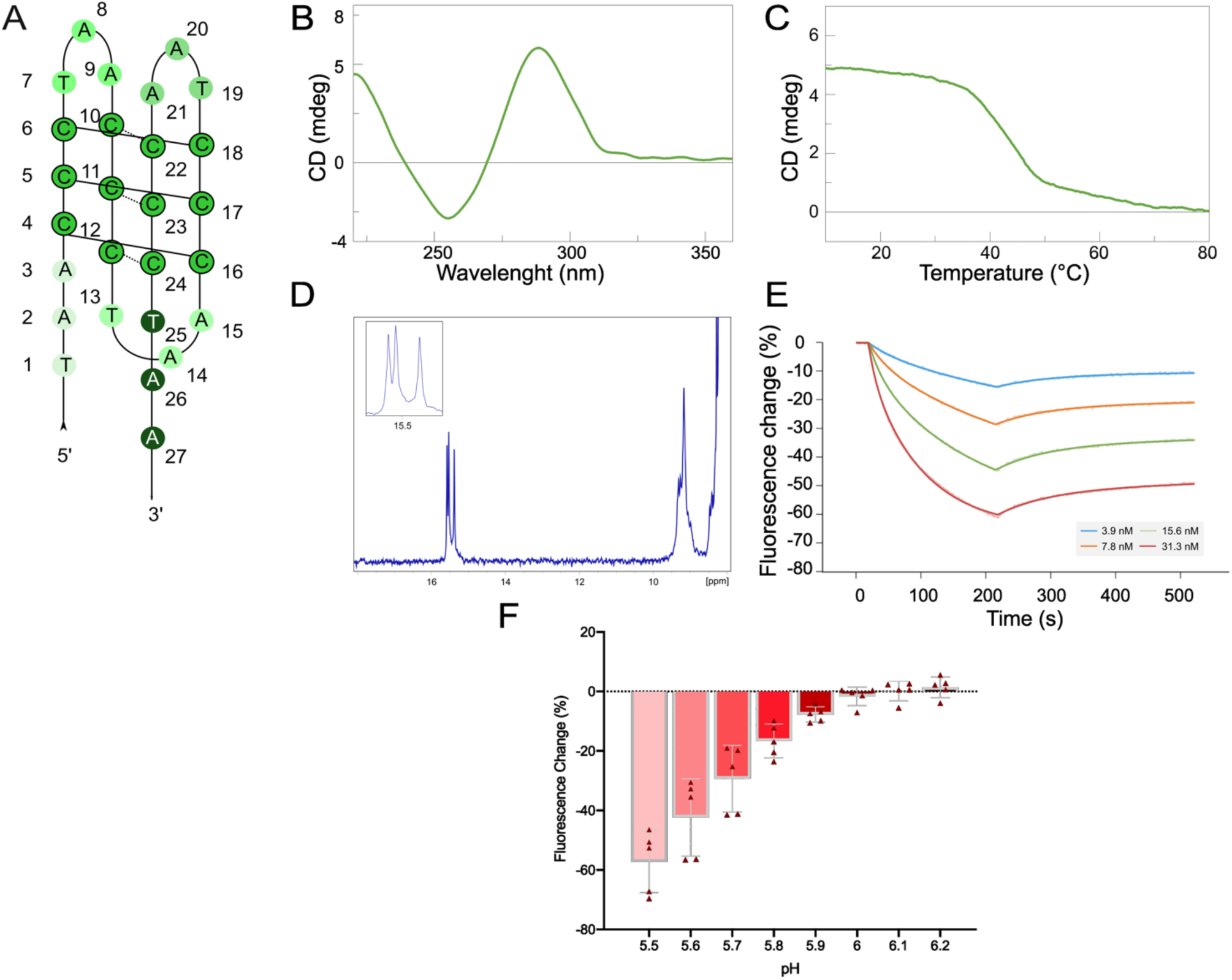
Biophysical characterization of the telomeric iM. A) Schematic representation of the C-NAT ODN containing iM. Numbers indicate each nucleotide position in the ODN. The cytosine core is monochromatically colored and hydrogen bonds are represented by a line between cytosines. B) CD spectrum of the C-NAT. On the x-axis, the wavelength is reported (nm), while on the y-axis, the CD value (mdeg). C) CD melting profile of the C-NAT. Temperature (expressed in °C) and CD (expressed in mdeg) are reported on the x- and y-axis, respectively. D) 1D ^1^H NMR spectrum of the C-NAT. E) Real-time fluorescence signals and fits of a representative experiment measuring the association and dissociation phases of iMab at stated concentrations towards immobilized C-NAT at pH 6.5. Time (expressed in s) and Fluorescence change (expressed in %) are reported on the x- and y-axis, respectively. F) Histogram showing the fluorescence change of C-NAT dye environment (expressed in %, y-axis) caused by each different pH (x-axis), calculated in the plateau phase of five different replicates.

### APE1 is able to stably bind iM telomeric sequences

Then we investigated APE1 ability to bind C-NAT *in vitro*. Firstly, we performed a SWB assay by analyzing the binding ability of equal amounts of recombinant purified APE1 wild-type (APE1^WT^) and its mutant lacking the first 33 residues (APE1^NΔ33^) (Fig. 2A and 2B and Supplementary Fig. S2A). The binding of both proteins toward C-NAT (Fig. 2A, lanes 3 and 4) was observed and resulted specific, since the inability to bind exhibited by BSA protein (Fig. 2A, lane 2). Although not statistically significant, we observed an almost 20% reduction in the binding ability by the APE1^NΔ33^ mutant compared to APE1^WT^, suggesting a role of the first 33 N-terminal amino acids in regulating the binding to the C-NAT. To deepen the observed difference, we employed the SwitchSENSE technology by functionalizing the chip surface with C-NAT and modulating the ligand concentration of both APE1^WT^ and APE1^NΔ33^ recombinant proteins at pH 5.5. By fitting the dose response curves in a monophasic model, we obtained similar KD for the two proteins, respectively of 6.48 nM (± 1.18) for APE1^WT^ and of 5.46 nM (± 1.38) for APE1^NΔ33^, demonstrating that both proteins show high affinity towards C-NAT with a minimal impact of the N-terminal region of APE1 in the binding affinity (Fig. 2C and Supplementary Fig. S2B and S2C). Additionally, 2D NMR spectroscopy analysis showed that the interaction between the C-NAT and APE1^WT^ caused the reduction of some peak intensities and chemical shift variations (Δδ) (Fig. 2D). The progressive decrease of peak intensities from the free protein (red) upon increasing concentrations of C-NAT (green and blue) suggested that the binding occurs in the intermediate exchange regime on the NMR timescale. The greater reduction in peak intensities at a 1:1 ratio (blue spectrum) suggested that a higher fraction of APE1^WT^ is engaged in binding, likely reaching near saturation. Moreover, as we can observe from Fig. 2E and Supplementary Fig. S2D, the C-NAT bound preferably to a distinct region of the canonical duplex DNA binding. The residues most involved in the binding are concentrated within and around α-helices 7 and 9. These helices are within the APE1 catalytic domain and contribute to DNA binding and stabilization of the enzyme-substrate complex. Interestingly, all these experiments were performed at pH 5.5, that contributed to Histidine (His) protonation and the increase of basic surfaces recognized by nucleic acids. In total, 4 His are within this new basic surface: H151 and H215 underwent changes, and H255 and 289 (blue) were not unambiguously identified in the 2D NMR spectra.

**Figure 2:**
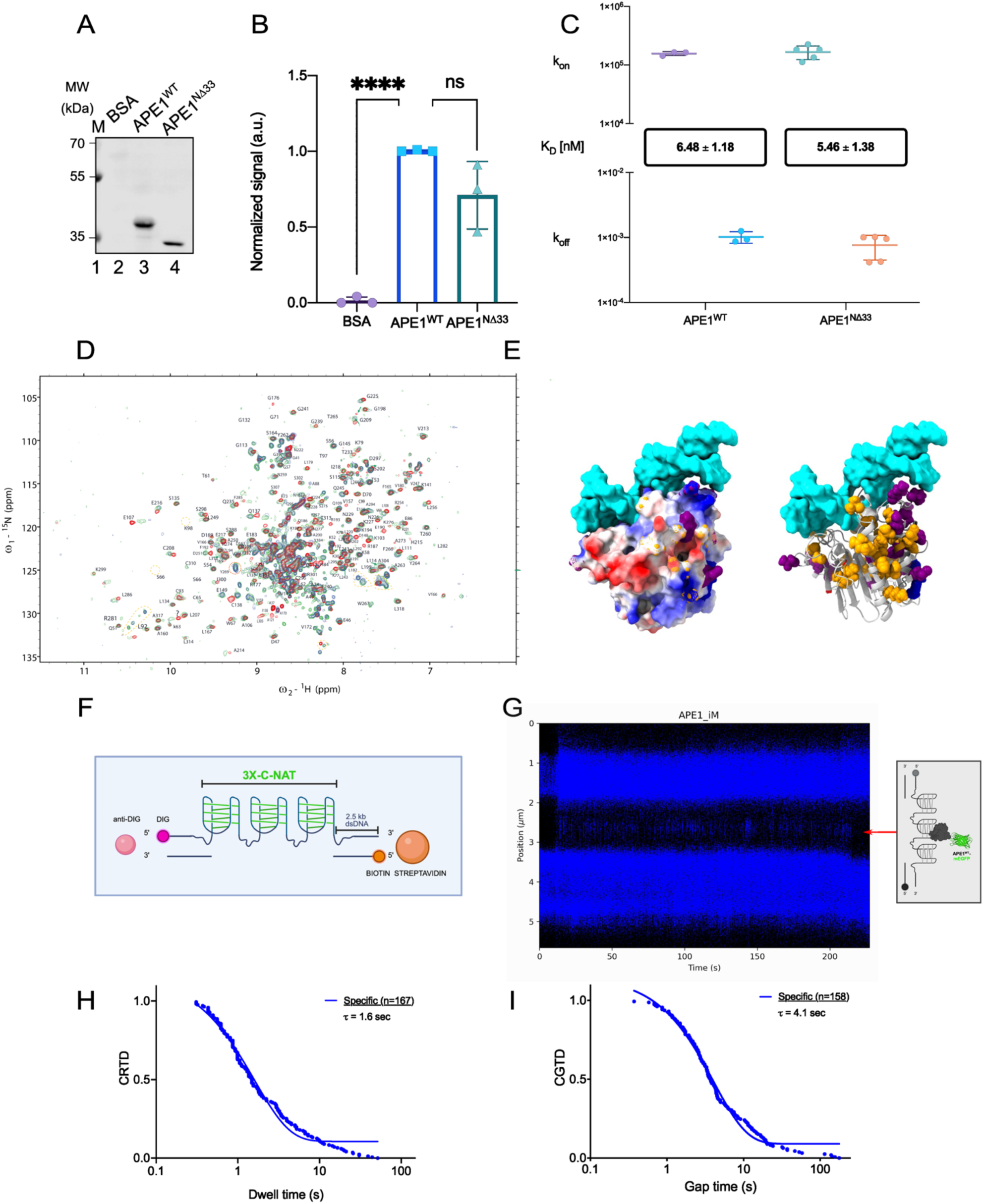
APE1 binds iM telomeric structures. A) Representative SWB assay between the C-NAT and recombinant APE1^WT^ and APE1^NΔ33^ proteins. BSA was used as a negative control. On the left, the electrophoretic marker is loaded, and the different molecular weights are expressed in kDa. B) Relative histogram summarizing the different binding abilities of three independent replicates towards C-NAT between BSA, APE1^WT^, APE1^NΔ33^, normalized on the respective total protein staining signal (Figure S2A). C) Graph representing the summary of C-NAT-APE1^WT^ and C-NAT-APE1^NΔ33^ kinetic parameters (kon and koff), expressed in nM, in MES buffer pH 5.5. KD values, expressed in nM, are reported (n=3). D) Representation of 2D {^15^N-^1^H} TROESY-HSQC NMR spectra recorded at 25 °C. Measurements were performed with APE1^WT^ alone (red), 0.6 (green) and 1:1 (blue) molar ratios of C-NAT to APE1^WT^. E) Depiction of NMR-derived binding analysis on APE1 structure (PDB code: 1bix). Cyan represents deposited canonical duplex DNA (for reference only). Purple represent residues which chemical shift variations (Δ8) that, before and after inclusion of i-motif DNA, are equal or superior to 2 standard deviations from the mean. Orange represent residues that lost over 2/3 or the respective peak volume. F) Schematic representation of the three consecutive telomeric iM (3x-C-NAT) embedded in a DNA sequence used for the single molecule analysis. G) Representative kymograph of APE1^WT^-mEGFP from A549 clone 31 NCE binding to the iM (3x-C-NAT) substrate. Time (expressed in s) and position (expressed in µm) are reported on the x- and y-axis, respectively. The iM position is indicated with the red arrow. H) Cumulative residence time distribution (CRTD) plot of specific binding events of APE1^WT^-mEGFP to 3x-C-NAT. I) Cumulative gap time distribution (CGTD) plot of APE1^WT^-mEGFP association with 3x-C-NAT under the tested conditions.

To characterize the binding of APE1 to the telomeric iM structure in a more physiological context, we performed a SMADNE analysis with the C-Trap^®^ technology (36). To perfom this experiment, we designed a DNA sequence containing three consecutive telomeric iMs (called 3x-C-NAT), ligated in the middle of a construct composed of two 2.5kb double-stranded DNA tracts on each side, called DNA handles, derivatized at their extremity with three biotins or three digoxigenins respectively (Fig. 2F). To set up the assay in the microfluidic system, we modulated the distance between the two beads, at pH 5.5, and detected the efficient formation of iM structures, which unfolded at an average force of 22 pN as evident from the force-distance (FD) curves of the tethered DNA (Supplementary Fig. S2E). Then we performed the SMADNE analysis by monitoring the iM binding of single mEGFP labelled APE1 proteins in the context of the nuclear cell extract. Specifically, we used NCE obtained from A549 clone 31 cell line, endogenously expressing APE1-mEGFP and deeply characterized in Supplementary Fig. S3. To visualize and asses APE1 binding to iMs, 3x-C-NAT DNA was kept at a force of 15 pN and exposed to the NCE (Fig. 2G). Importantly, we observed numerous specific binding events of APE1-mEGFP to the iM. Indeed, the duration of the iM binding events had an average lifetime (𝜏) of 1.6 seconds (Figure 2H) with 4.1 second gap times between the binding events (Figure 2I). All these findings clearly confirmed the ability of APE1 to specifically bind the C-NAT sequence and suggested that the unstructured N-terminal region does not affect the overall affinity of the complex but rather plays a significant role in stabilizing the DNA-protein complex formation.

### The position of AP-sites in the iM influences its stability and APE1 binding

APE1 repair efficiency of AP-sites embedded in iM tetraplex structures is unknown and requires investigation. We aimed to characterize the cleavage activity of APE1 on damaged iM by evaluating its endonuclease activity on AP-sites placed at different positions in the C-NAT sequence. Specifically, we designed: AP14 and AP20 ODNs bearing an AP-site in the second and in the third loop, respectively, both substituting an adenine base; AP16 and AP17 ODNs holding the AP-site in the core of the iM structure, substituting the first and middle cytosine of the third C-run, respectively (Fig. 3A and Table 1).

**Figure 3:**
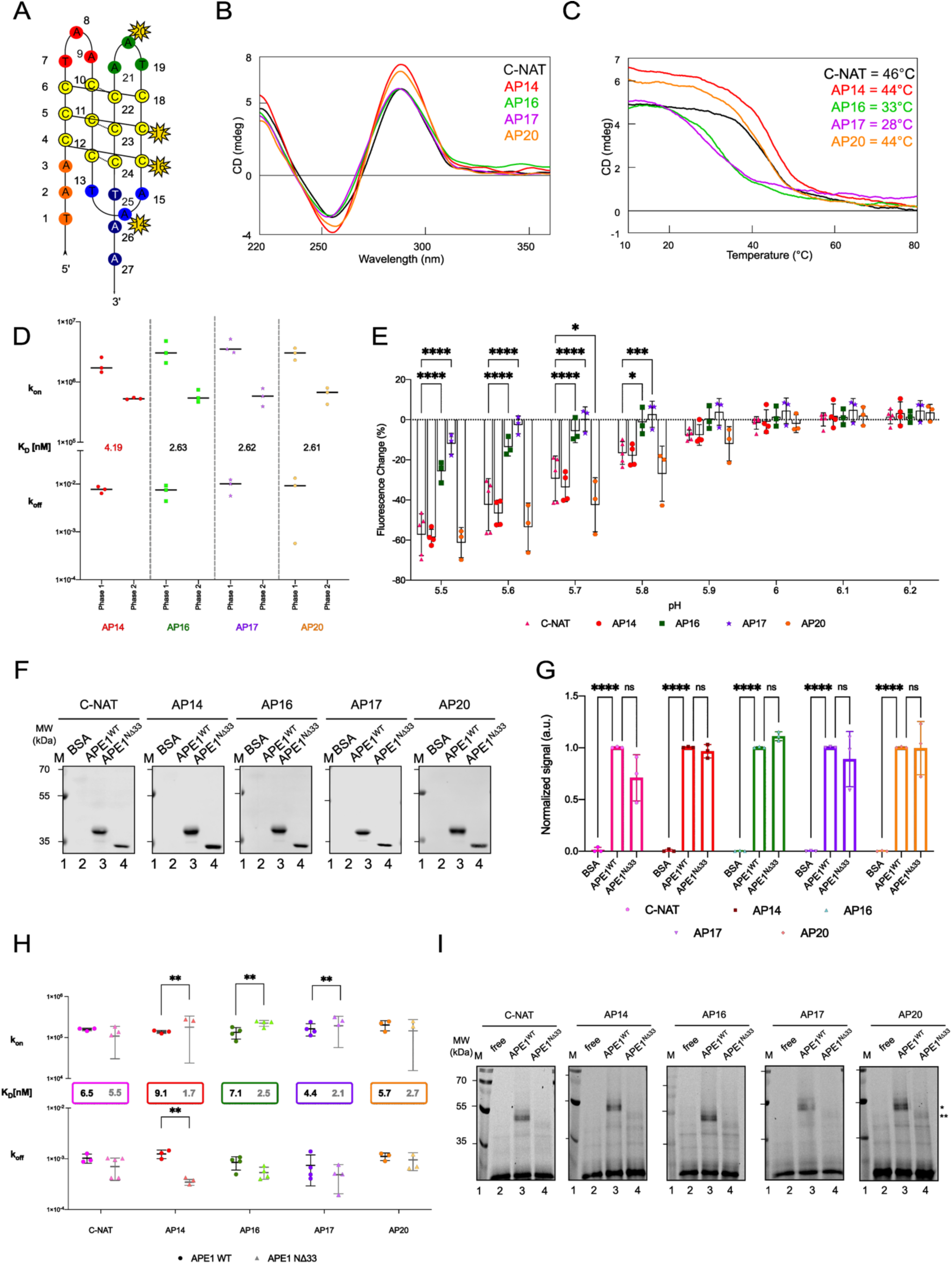
iM are structurally affected when containing AP-sites and are specifically bound by APE1. A) Scheme of the damaged telomeric iM ODNs, holding an abasic site at different positions (AP14, AP16, AP17, AP20), indicated by a star. B) CD spectrum of damaged ODNs. On the x-axis, the wavelength is reported (nm), while on the y-axis the CD value (mdeg). C) CD melting profile of damaged ODNs. Temperature (expressed in °C) and CD (expressed in mdeg) are reported on the x-and y-axis, respectively. D) Graph representing the summary of iMab kinetic parameters, including Kon and Koff, towards each of the ODNs examined in MES buffer pH 5.5. KD values, expressed in nM, are reported (n=3). E) Histogram showing fluorescence change of the different ODNs dye environment (expressed in %, y-axis) caused by increasing pH (x-axis), calculated in the plateau phase of five different replicates. F) Representative SWB assays between each ODNs, as indicated upon each panel, and APE1^WT^ or APE1^NΔ33^ proteins. BSA was used as a negative control. On the left, the electrophoretic marker is loaded, and the different molecular weights are expressed in kDa. G) Relative histogram summarizing the different binding abilities of three independent replicates towards the ODNs used in this study between BSA, APE1^WT^, APE1^NΔ33^, normalized on the respective total protein staining signal (Figure S4C). H) Graph representing the summary of APE1^WT^ and APE1^NΔ33^ kinetic parameters, including kon and koff, towards each of the ODNs examined in MES buffer pH 5.5. KD values, expressed in nM, are reported (n=3). I) Representative crosslinking analysis with APE1^WT^ and APE1^NΔ33^ recombinant proteins and each damaged iM (25 nM). On the left, the electrophoretic marker is loaded, and the different molecular weights are expressed in kDa.

Firstly, we assessed the impact of AP-sites on the formation and stability of the iM itself through CD spectroscopy. As shown in Fig. 3B, all the AP-site-containing sequences showed CD profiles similar to that of the C-NAT with a positive band at 288 nm and a negative one at 255 nm (Fig. 1C). Specifically, AP14 and AP20 had a Tm of 44°C, comparable to the C-NAT (46°C), while AP16 and AP17 of 33°C and 28°C, respectively (Fig. 3C), suggesting that iMs formed by AP16 and AP17, which present AP-sites in the C-core, are much less stable with respect to those having these sites in the loops, AP14 and AP20. We also used the two SwitchSENSE approaches, previously described, to further analyze potential differences in the iM folding ability among the damaged sequences. We firstly validated the iM folding with the iMab antibody at pH 6.5 (Fig. 3D and Supplementary Fig. S4A). The best-suited fitting algorithm was the biphasic model, in which only the first dissociation constant (KD1|1) of the iMab towards the different ODNs was calculable. Despite the different position of AP-sites within the iM, iMab bound each ODN with dissociation constants in the low nanomolar range, with minor differences between them. Furthermore, by using the second approach (Fig. 3E and Supplementary Fig. S4B), we found that AP14 and AP20 have a similar behaviour to the C-NAT, showing parallel levels of fluorescence change (%), whereas AP16 and AP17 behave in a significantly different way since, already at pH 5.5, they showed a 35% difference in fluorescence change compared to the undamaged C-NAT, indicating that a lower percentage of the total amount of these ODNs was folded compared to the undamaged one. Moreover, in agreement with the Tm values, we observed a difference between AP16 and AP17, as AP16 seemed to be slightly more stable than AP17. In summary, whereas C-NAT, AP14 and AP20 were stable until reaching pH 6.0, AP16 and AP17 unfolded at pH 5.8 and 5.7, respectively, thus confirming that the position of the AP-site crucially influences iM structure.

Subsequently, the ability of APE1 to recognize damaged telomeric iM structures was investigated *in vitro*: in SWB assay, by using equal amounts of APE1^WT^ and APE1^NΔ33^ (Supplementary Fig. S4C), both proteins bound modified iM with not statistically significant differences (Fig. 3F and 3G). The SwitchSENSE assay was also employed and, by fitting the dose response curves with a monophasic model, we obtained KD values in the nanomolar range for both proteins for each ODN. In detail, APE1^WT^ bound the ODNs with similar affinities (Fig. 3H and Supplementary Fig. S4D), while APE1^NΔ33^ exhibited higher affinity towards the damaged ODNs compared to the C-NAT (Fig. 3H and Supplementary Fig. S4E), and also in comparison with APE1^WT^. To deepen these evidences, we employed a UV-crosslinking coupled to SDS-PAGE analysis (Fig. 3I). We observed that APE1 was able to bind each ODN forming a stable complex of 50 kDa (as indicated by a single asterisk) while for APE1^NΔ33^ weak bands were detected, as indicated by a double asterisk, suggesting that the N-terminal region of APE1 can concurr to stabilize the complex without affecting the overall binding affinity.

### APE1 endonuclease activity on iM strongly depends on AP-site position and on its unstructured N-terminal region

To test if the AP-site position on the telomeric iM structure might influence APE1 endonuclease activity, we performed an AP-site incision assay (Fig. 4A). As a positive control of APE1^WT^ activity, we used a single-stranded oligonucleotide (ss_dF, Table 1), designed to lack any secondary structure formation ability. As expected, ss_dF was the most efficiently processed, reaching a 66% of cleavage after 60 minutes of incubation (Fig. 4B). Interestingly, AP16 and AP17 were preferentially cleaved with respect to AP14 and AP20, reaching ∼25% of cleavage activity by APE1^WT^ upon 60 minutes. On the contrary, AP14 reached ∼10% of cleavage, whereas AP20 was completely not cleaved (Fig.4B): these data pointed out that APE1^WT^ cleaves more efficiently damaged iM bearing the AP-site in the core of the structure rather than in the loops and that these differences could be connected to the higher ability of AP14 and AP20 to conserve iM folding with greater stability, which presumably inhibits APE1^WT^ endonuclease activity. To speculate this aspect, we performed the cleavage assay by using the ODNs in a regular Watson-Crick double-stranded conformation in order to avoid iM structure folding (Table 1). As shown in Supplementary Fig. S5A, APE1^WT^ was able to cleave each ds_ODN at a lower concentration than the ss_ODN counterpart (0.125 nM *versus* 80 nM, respectively) and more rapidly (15 minutes *versus* 60 minutes, respectively). Indeed, at 15 minutes, both ds_AP14 and ds_AP16 reached a ∼50% of cleavage, while both ds_AP17 and ds_AP20 reached the ∼30% (Supplementary Fig. S5B-E), corroborating previous data showing that AP-sites located towards the 3’ were processed slightly less efficiently in the dsODNs but at the same time confirming a major effect of the AP-site position within the iM structure.

**Figure 4:**
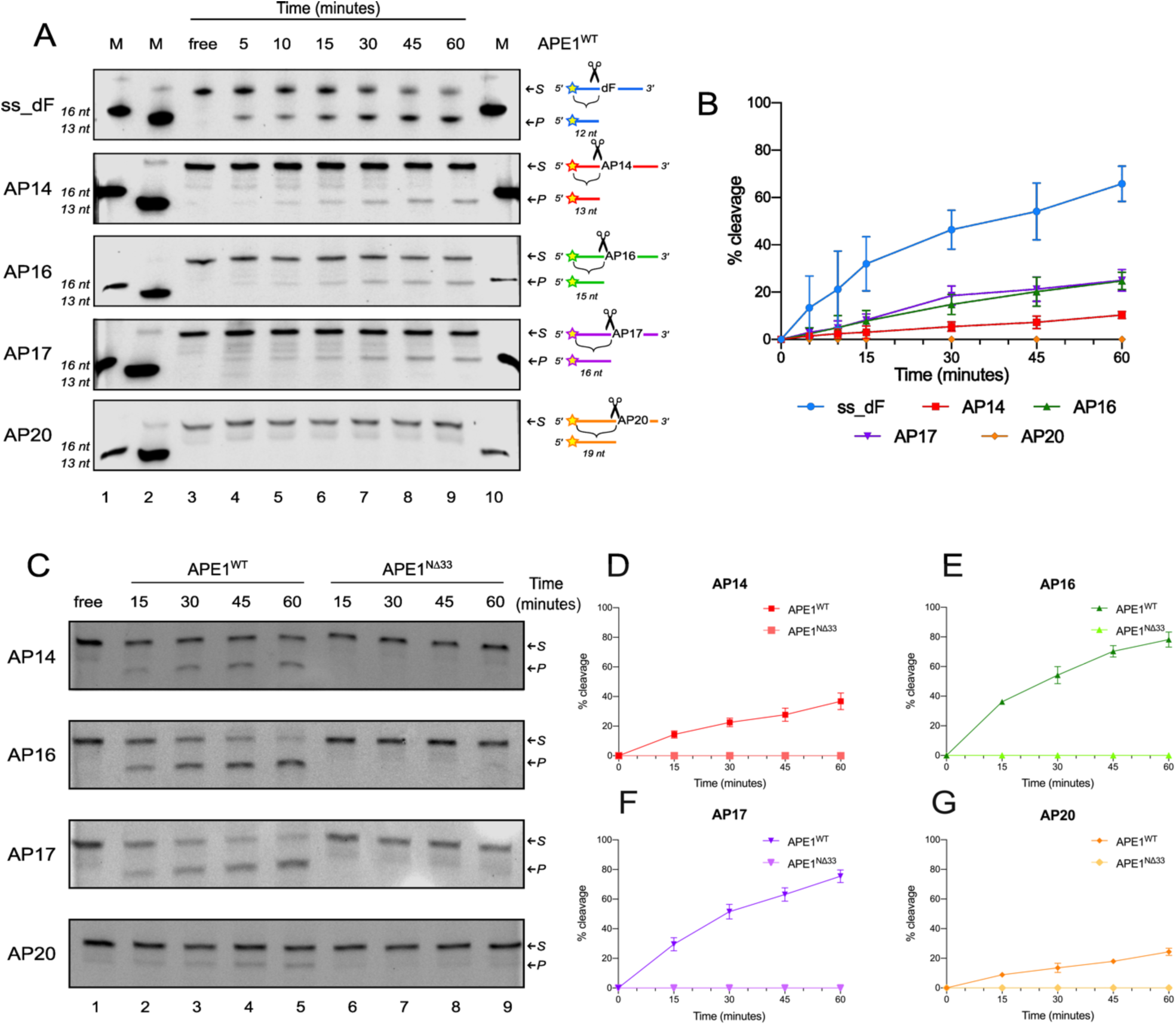
The position of the AP-site on the telomeric iM influences APE1 endonuclease activity, which relies on its N-terminal region. A) Representative denaturing polyacrylamide gels of time-course kinetics APE1^WT^ cleavage activity. Lanes 1 and 2 indicate DNA fragments with known length (16 and 13 nucleotides, respectively). The “free” sample represents the control without protein (lane 3). On the right, the substrate and the product bands are indicated by two arrows and the expected length of the cleavage product is indicated upon the AP-site. A constant dose of APE1 (80 nM) was incubated with the respective oligonucleotide at 37°C. B) Relative graph depicting the time (expressed in minutes) and percentage of cleavage (%) on the x- and y-axis, respectively. Data are expressed as mean ± SD of three independent technical replicates. C) Representative denaturing polyacrylamide gels of APE1^WT^ (lane 2-5) and APE1^NΔ33^ (lane 6-9) cleavage activity performed on all substrates (lane 2-9). “free” sample represents the control without protein (lane 1). On the right, the substrate and the product bands are indicated by two arrows. A constant dose of APE1^WT^ or APE1^NΔ33^ (120 nM) was incubated with the respective oligonucleotide at 37°C, and the reactions were stopped at different time points, indicated upon the gel. D-G) Relative graphs illustrating the time-course kinetics activity of APE1^WT^ and APE1^NΔ33^ recombinant proteins on AP14 (D), AP16 (E), AP17 (F) and AP20 (G). Time (expressed in minutes) and percentage of cleavage (%) are reported on the x- and y-axis, respectively. Data are expressed as mean ± SD of three independent technical replicates.

In order to evaluate the impact of the absence of the N-terminal region of the protein on its catalytic activity on damaged iM, we performed similar epxeriments by employing APE1^NΔ33^ and surprisingly found that APE1^NΔ33^ did not cleave any substrate (Fig. 4C-G). As control, the cleavage assay by using ds_ODNs was carried out (Supplementary Fig. S5): in this case APE1^WT^ and APE1^NΔ33^ processed each ds_ODN to a comparable extent, as already reported (41). These data clearly indicate a stabilizing role in binding the substrate played by the APE1 N-terminal region as essential for its endonuclease activity on lesions embedded on alternative DNA secondary structures as G4 and iM.

### APE1 enzymatic activity on damaged telomeric iM is inhibited by Poly(rC)-binding protein 1

In our previous studies on APE1-interactome in HeLa cells (42), we found several human ribonucleoproteins (hnRNP), which are involved in modulating iM folding, including PCBP1 (23). Indeed, PCBP1 binds to several iM, maintains their folding and plays a role in regulating the competing formation of iM and G4 in the genome of human cells (23). Therefore, we better characterize the close proximity (<40 nm) between APE1 and PCBP1 in our telomeric model, using PLA analysis By using PLA analysis, the occurrence of APE1-PCBP1 interaction in HeLa cells was confirmed (Fig. 5A and Supplementary Fig. S6A) and evaluated also in U2OS cells from osteosarcoma (Fig. 5A and Supplementary Fig. S6B). APE1 specifically interacted with PCBP1, in both nuclear and cytosolic compartments, with a higher number of PLA dots in the nucleus with a mean of 70 dots *per* nucleus in HeLa and 84 dots *per* nucleus in U2OS (Fig. 5B). This data was also confirmed by PLA analysis on HeLa cell transiently silenced for APE1, PCBP1 or both proteins, leading to the loss of the interaction (Supplementary Fig. S6C), as well as by analyzing two stable APE1 knock-out (KO) clones, derived from U2OS cell line, called U2OS^19^ and U2OS^21^, which were transiently silenced for PCBP1 (Supplementary Fig. S6D). In both models, the deficiency of one or both proteins resulted in the loss of PLA dots, as expected.

**Figure 5:**
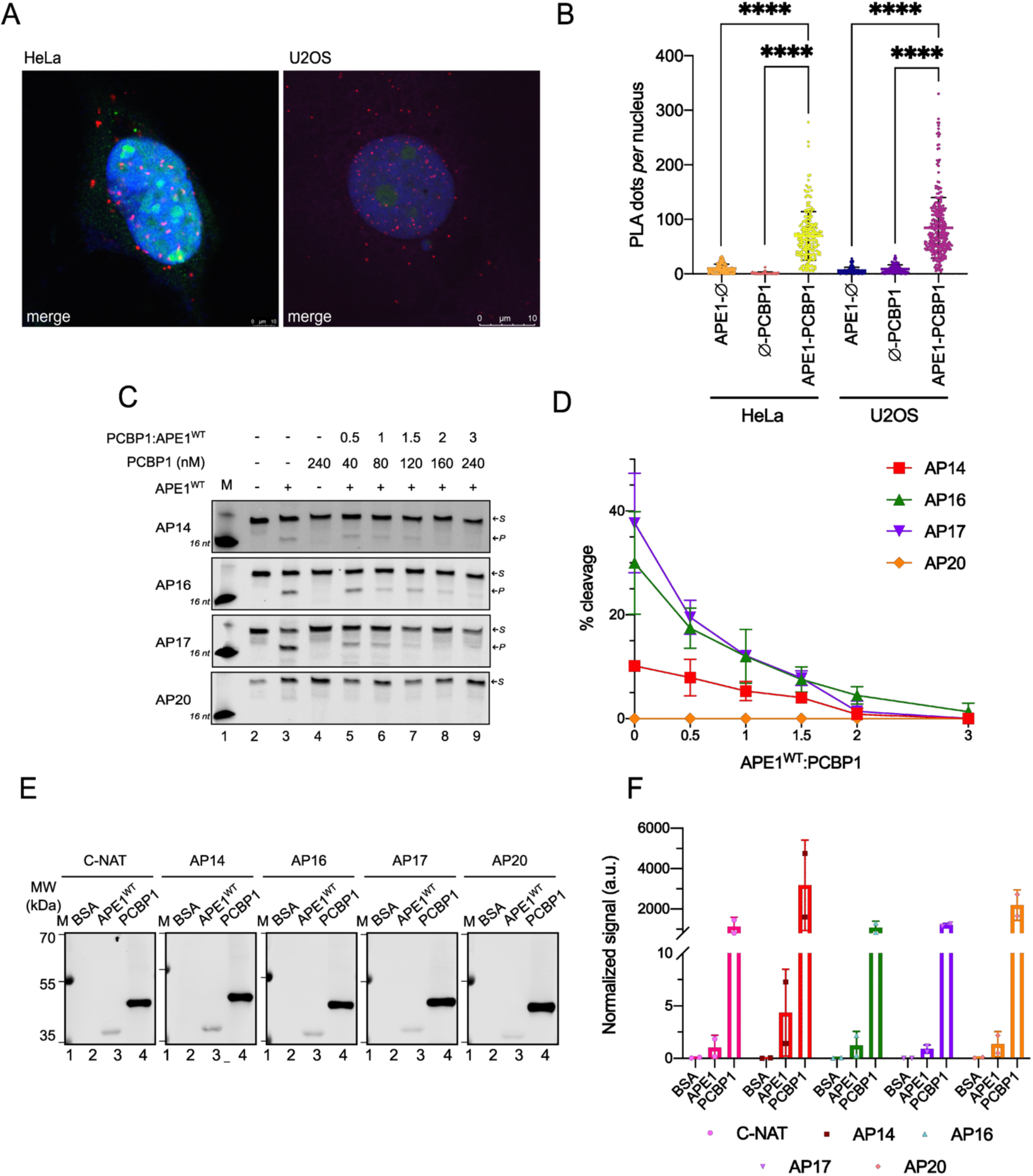
Damaged telomeric iM cleavage mediated by APE1 is modulated by APE1 interaction with PCBP1. A) PLA analysis of the interaction between APE1 and PCBP1 proteins in HeLa and U2OS cells (PLA dots in red). The merged panel shows the overlay between the four channels, including APE1 and PCBP1 staining in green (rabbit-488) and magenta (mouse-633), respectively, and the nuclear DAPI staining in blue. The scale is indicated and expressed in μm. B) Graph depicting the number of PLA dots *per* nucleus in U2OS and HeLa cell lines. Average and standard deviation values are plotted (n=3). C) Representative denaturing polyacrylamide gels of APE1^WT^ cleavage activity performed on all substrates (lane 3, 5-7), modulated by the co-incubation with PCBP1 (lane 5-7). Lane 2 is a control without any proteins. Lane 4 is a control with only the substrate and PCBP1. Lane 1 indicates a DNA fragment with a known length (16 nt). On the right, the substrate and the product bands are indicated by two arrows. The ODNs were pre-incubated with increasing amounts of PCBP1 (indicated upon the gel and expressed in nM) for 90 minutes at 4°C. A constant dose of APE1^WT^ (80 nM) was then added to the reactions and incubated at 37°C for 60 minutes. D) Relative graph illustrating the activity of APE1^WT^ recombinant protein after pre-incubation with different amounts of PCBP1 on AP14, AP16, AP17 and AP20. PCBP1 concentration (nM) and percentage of cleavage (%) are reported on the x- and y-axis, respectively. Data are expressed as mean ± SD of three independent technical replicates. E) Representative SWB shows the binding between the telomeric iM ODNs (reported upon the gel) and recombinant APE1^WT^ and PCBP1 proteins. BSA was used as a negative control. Each protein was loaded on the SDS-PAGE gel, blotted on the membrane and then incubated with the respective fluorescent probe (5 pmol). On the left, the electrophoretic marker is loaded, and the different molecular weights are expressed in kDa. F) Relative histogram summarizing the different binding abilities towards each ODN between BSA, APE1^WT^, PCBP1 (n=3), normalized on the respective total protein staining signal (Figure S6G).

Then we investigated whether PCBP1 exerts its role in maintaining genome integrity (23) by functionally modulating the enzymatic activity of APE1 on damaged iM structures. As expected, PCBP1 was not able to exert any endonuclease activity on damaged iMs alone (Fig. 5C, lanes 4). Interestingly, the pre-incubation of damaged iM with PCBP1 and the subsequent addition of APE1^WT^ had a strong dose-dependent inhibitory effect on APE1^WT^ endonuclease activity, commonly observed for any ODNs (Fig. 5C, lanes from 5 to 9). Specifically, the pre-incubation with an under-stoichiometric dose of PCBP1 compared to APE1^WT^ (ratio: 0.5:1 – lane 5) was already sufficient to inhibit the product formation of ∼50% for AP16 and AP17, while of about 23% in the case of AP14. As expected, AP20 that is not efficiently processed by APE1^WT^, was not influenced by PCBP1 presence (Fig. 5D). By performing the same experiment on the ds_ ODNs (Supplementary Fig. S6E and S6F), we observed that the pre-incubation with PCBP1 did not influence the cleavage ability of APE1^WT^, confirming that the inhibitory effect is specific for the presence of iM structure. Lastly, to assess if the inhibitory effect of PCBP1 was due to a possible competition with APE1 for the iM, we performed a SWB assay by loading equal amounts of BSA (as negative control) and APE1^WT^, together with 40-fold lower amount of PCBP1 protein (Fig. 5E). After signal normalization on total protein staining (Supplementary Fig. S5G), we observed that the binding of PCBP1 was at least 1000-fold higher compared to that of APE1^WT^ for each ODN (Fig. 5F). Overall, these results suggested that PCBP1 has a higher binding affinity for the iM telomeric compared to APE1. Moreover, PCBP1 could shield these structures, exerting a competitive effect on APE1, thus inhibiting its cleavage activity.

### APE1 and PCBP1 depletion impacts on telomere length and on the interaction with Telomeric repeat-binding factor 2

To further explore the biological relevance of both APE1 and PCBP1 proteins, we examined their impact on telomere length maintenance by using U2OS cell line and their corresponding APE1-KO models. Due to the great length of U2OS telomeres, previous studies unsuccessfully tried to determine size changes upon transient APE1 silencing by using TRF assay (43). In this context, we performed TRF assay (Fig. 6A) by sonicating the DNA obtained from U2OS^WT^ and U2OS^19^ cell lines, transiently silenced for PCBP1 (Supplementary Fig. S7A), before digesting with restriction enzymes (Supplementary Fig. S7B). Interestingly, APE1 KO led to a slight but significant increase in telomere length (Fig. 6B), whereas the transient silencing of PCBP1 in both WT and KO models led to the same slight but significant decrease of telomere length (Fig. 6C). The opposite behaviour of these two proteins could be explained by their roles in maintaining telomere length at equilibrium when both proteins are present.

**Figure 6:**
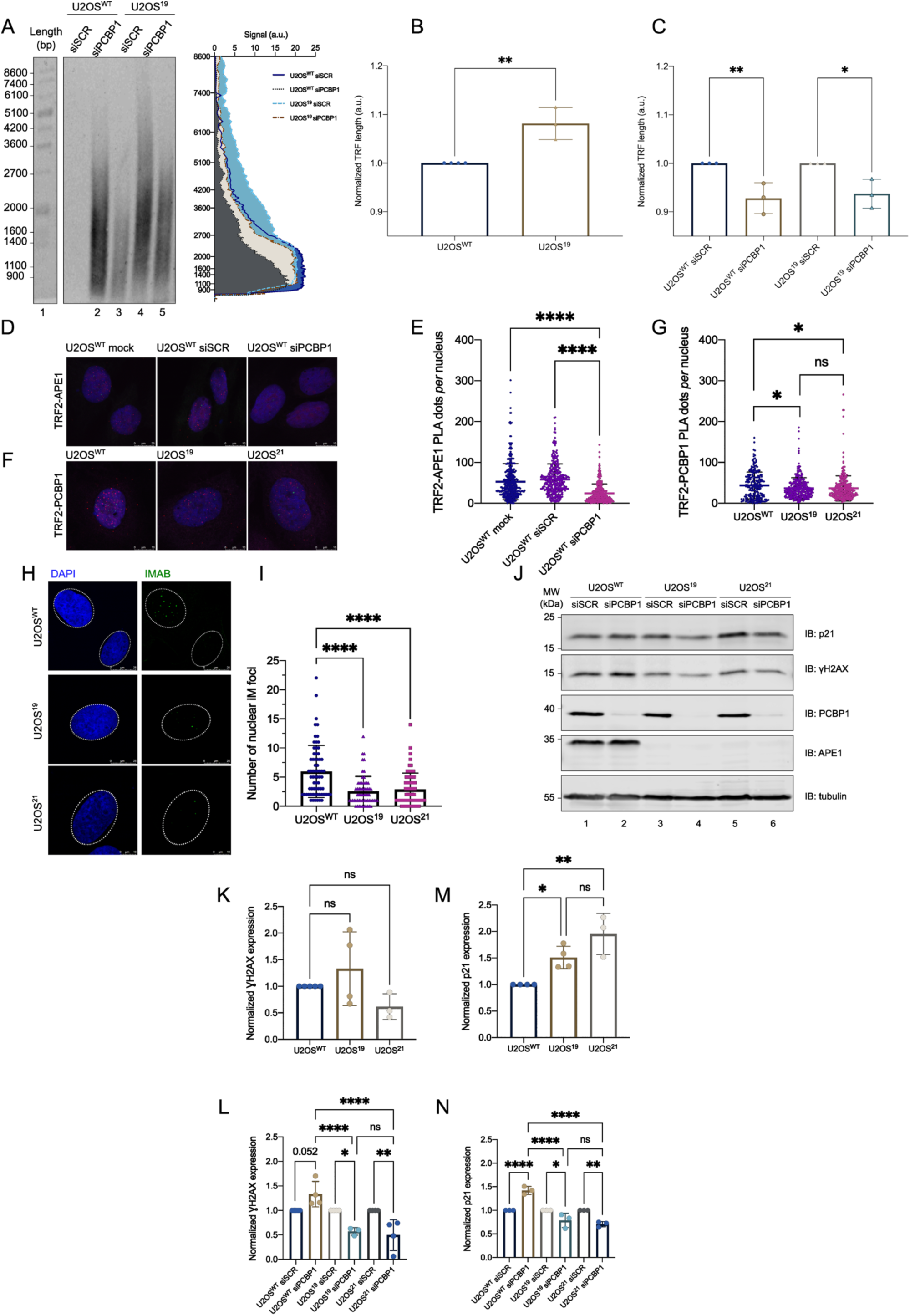
APE1 and PCBP1 depletion impacts on telomere length and on the interaction with TRF2 protein. A) TRF assay was used to measure telomere length in U2OS cells expressing (U2OS^WT^) or knocked-out for APE1 (U2OS^19^) protein, either silenced or not for PCBP1 protein (siPCBP1 and siSCR, respectively), as indicated upon the gel. On the left side of the gel, the molecular weight marker, as provided by the kit, is loaded, and the length of each band is indicated and expressed as bp. On the right side, a representative graph showing the intensity profiles relative to the TRF assay is reported. On the y-axis, the signal intensity is reported (expressed in a.u.), while on the x-axis the TRF marker length (expressed in bp) is reported. B) Graph reporting the normalized TRF mean length of U2OS^WT^ and U2OS^19^. TRF mean length is normalized to U2OS^WT^ values. Data are expressed as mean± SD of three independent replicates. C) Graph reporting the normalized TRF mean length of U2OS^WT^ and U2OS^19^ silenced for PCBP1. TRF mean length is normalized to each respective scramble silencing values. Data are expressed as mean± SD of three independent replicates. D) PLA analysis between TRF2 and APE1 proteins in U2OS cells, either silenced or not for PCBP1 protein (siPCBP1 and mock, siSCR, respectively). The representative merge panel shows PLA dots in red, TRF2 staining in green and APE1 staining in magenta. The nuclear staining, obtained with DAPI, is in blue. The scale is indicated and expressed in μm. E) Relative graph depicting the number of TRF2-APE1 PLA dots *per* nucleus in all U2OS^WT^ conditions. Average and standard deviation values are plotted (n = 3). F) PLA analysis of the interaction between TRF2 and PCBP1 proteins in U2OS cells expressing (U2OS^WT^) or knocked-out for APE1. The representative merge panel shows PLA dots in red, TRF2 staining in green and and PCBP1 in magenta. The nuclear staining, obtained with DAPI, is in blue. The scale is indicated and expressed in μm. G) Relative graph depicting the number of TRF2-PCBP1 PLA dots *per* nucleus in U2OS^WT^, U2OS^19^ and U2OS^21^ cell lines. Average and standard deviation values are plotted (n = 3). H) Representative immunofluorescence images of iMab antibody staining in U2OS^WT^, U2OS^19^ and U2OS^21^ cell lines. iMab foci are shown in green (rabbit 488 secondary antibody), while nuclei were stained with DAPI. The scale is indicated and expressed in μm. I) Scatter plot of iMab analysis representing individual values of iM foci in the nuclear compartment of U2OS clones. Foci were counted for each cell, using DAPI as a nuclear mask. Data are expressed as means ± SD of two independent replicates. J) Western blot analysis of p21, 𝛾H2AX, PCBP1 and APE1 levels in U2OS^WT^, U2OS^19^ and U2OS^21^ cell lines, either silenced or not for PCBP1 (siPCBP1 and siSCR, respectively). Tubulin was used as normalizer. Molecular weights (MW, expressed in kDa) are reported on the left. K-L) Densitometric analysis of 𝛾H2AX expression levels, normalized to tubulin. In the upper panel (K), fold change values relative to U2OS^WT^, arbitrarily set to 1, are shown. In the lower panel (L), fold change values relative to each siSCR, arbitrarily set to 1, are shown. Values are mean ± SD of four independent replicates. M-N) Densitometric analysis of p21 expression levels, normalized to tubulin. In the upper panel (M), fold change values relative to U2OS^WT^, arbitrarily set to 1, are shown. In the lower panel (N), fold change values relative to each siSCR, arbitrarily set to 1, are shown. Values are mean ± SD of four independent replicates.

Previous studies have observed that both APE1 and PCBP1 could have a role in the maintenance of telomeres through the regulation of proteins belonging to the shelterin complex. Indeed, APE1 co-localizes with both TRF2 and POT1 proteins (43), whereas PCBP1 co-localizes with TRF2 and Telomeric Repeat-binding Factor 1 (TRF1) (14). TRF1 and TRF2 bind the double-stranded region of telomeres and, specifically, TRF2 prevents the recognition of telomere ends as double-strand breaks, inhibiting Ataxia Telangiectasia Mutated (ATM)-activated DDR (44). POT1 binds the G-rich overhang and protects it from cleavage (44). The displacement of TRF2 and POT1 from telomeres is an important sign of telomere dysfunction (44). It has been demonstrated that the depletion of APE1 leads to a decreased amount of TRF2 at telomeres, without affecting the overall amount of TRF2 in the cell (43). To better study APE1 and PCBP1 role and their interplay at the telomere level, we examined their interaction with TRF2. Specifically, we firstly employed PLA analysis between APE1-TRF2 (Supplementary Fig. S7C-D) and PCBP1-TRF2 (Supplementary Fig. S7E-F) in U2OS^WT^ cells. Both APE1 and PCBP1 interacted with TRF2 in the nuclear compartment, with a mean of 53 dots *per* nucleus for APE1-TRF2 (Supplementary Fig. S7D) and of 43 dots *per* nucleus for PCBP1-TRF2 (Supplementary Fig. S7F). In order to assess the existence of any influence between APE1 and PCBP1 in the modulation of their interaction with TRF2, we transiently silenced PCBP1 in U2OS^WT^ cells and we detected the presence of APE1-TRF2 PLA dots (Fig. 6D and Supplementary Fig. S7G). Although APE1 staining (in magenta) and TRF2 foci (in green) were both well-detectable after silencing PCBP1 without any effect, the PLA dots were far less apparent compared to the mock and siSCR conditions, with a dramatic reduction of about 60% compared to the siSCR condition (Fig. 6E). At the same time, we evaluated the interaction of PCBP1-TRF2 in both U2OS^19^ and U2OS^21^ APE1-KO cell models (Fig. 6F and Supplementary Fig. S7H). Again, although PCBP1 and TRF2 staining was not affected by the absence of APE1, the PLA dots upon APE1 KO were less and more faint compared to the WT, with a 16% reduction (Fig. 6G). In both PCBP1 silencing or APE1 KO conditions, the TRF2 expression levels evaluated by Western blot analysis didn’t show any significant downregulation (Figures S6I-J), confirming already previously published data (43).

Moreover, to further assess the interplay occurring between iM dynamics and APE1 in a cellular context, we explored whether APE1 depletion could influence the accumulation of iM foci in the nuclei of U2OS cells. With this aim, we performed an immunofluorescence analysis by using iMab antibody (Fig. 6H). The signal appears as a single discrete foci that can be quantified. Surprisingly, a significant depletion of iMab dots of about 50% was observed in both APE1 KO clones U2OS^19^ and U2OS^21^, thus connecting APE1 to iM folding regulation also in cells (Fig. 6I).

Lastly, we focused our attention on how APE1 and PCBP1 depletion could influence the DNA damage response pathway. Therefore, we analyzed the accumulation of DNA double-strand breaks by measuring the activation of 𝛾H2AX and a downstream target of the DDR and cell senescence, p21, which can be modulated at multiple levels by APE1 through its interplay with p53, upon depletion of either or both APE1 and PCBP1 proteins (Fig. 6J). APE1 depletion in U2OS^19^ and U2OS^21^ cell models did not result to a significant activation of 𝛾H2AX compared to U2OS^WT^ under basal conditions (Fig. 6K). On the contrary, PCBP1 depletion led to ∼35% increase in phosphorylated 𝛾H2AX (Fig. 6L), confirming previous results obtained in other cell lines (23). Surprisingly, when both proteins were depleted, phosphorylated 𝛾H2AX levels were even lower than in basal condition, specifically of the 40% for U2OS^19^ and 50% for U2OS^21^ compared to the respective siSCR (Fig. 6L), suggesting a possible impairment in the signaling of DNA damage when both proteins are absent. The levels of p21 protein after APE1 depletion in U2OS cell models resulted increased of the 50% in U2OS^19^ and 70% in U2OS^21^, confirming previous results linking APE1 depletion with p21 overexpression (45–47) (Fig. 6M). p21 levels were also higher upon PCBP1 silencing compared to siSCR, establishing that the absence of PCBP1 induces higher levels of DNA damage with downstream relevance to the activation of p21 (Fig. 6N). Lastly, we observed that p21 levels decreased when PCBP1 was silenced in U2OS^19^ and U2OS^21^, respectively, by the 22% in U2OS^19^ (compared to its siSCR) and of the 30% in U2OS^21^(compared to its siSCR), almost taking back p21 expression to its basal levels (Fig. 6N).

In summary, all these experiments allowed us to conclude that APE1 and PCBP1 contribute to telomere stability in ALT cells by maintaining telomere length at equilibrium when both present and by modulating the interaction of the other with shelterin protein TRF2.

## DISCUSSION

How genome stability maintainance is handled on DNA secondary structures is an emerging intriguing topic. To date, extensive work has been carried out to examine how BER proteins interact and repair G4 structures at promoters and telomeres, while other DNA secondary structures have been overlooked. In general, AP-site lesions embedded in G4 are repaired with really low efficiency compared to duplex DNA, leading to a hypothesis that links DNA damage with possible novel roles when localized at these specific regions. For example, oxidative modifications on G4-forming sequences are hypothesized to work as epigenetic regulators for modulating transcription (48).

Oxidation on guanines embedded in duplex G-rich DNA are handled by 8-oxoguanine glycosylase 1 (OGG1), generating AP-sites. The recognition of these AP-sites by APE1 stimulates the folding of the G4, acting as a transcriptional hub and modulating gene expression. Preliminary work by Burrows and Fleming group proposed that 8-oxoguanine modification can have analogous regulatory roles also when embedded in other kind of secondary structures (i.e. Z-DNA, cruciform DNA, iM), proposing a similar mechanism of action at promoter levels (49, 50). Up to now, DNA iM damage repair is still a completely underexplored field. The present study points to fill this knowledge gap by characterizing the effects of AP-sites on the folding of telomeric iM structures and on the binding and processing activity of APE1 on these damaged structures. Using several complementary biochemical, biophysical, structural and cellular analyses, we characterized the possible role of accessory regulatory proteins, such as PCBP1, in the modulation of APE1 enzymatic activity on iM telomeric structures and on their overall functional roles on maintenance of telomere stability.

Firstly, we examined C-NAT ODN iM folding ability. CD spectroscopy showed that the studied ODN folds in an iM at pH 5.5. Indeed, differences in the spectrum between the unfolded cytosine-rich strand and the folded iM are marked by an increase of the negative CD band from 245 nm to 260 nm, with a simultaneous decrease of the positive one from 288 nm to 275 nm (39, 40). This result was furthermore confirmed by 1D NMR, that could detect the typical imino-peaks around 15-16 ppm. While the iMab SwitchSENSE approach revealed that the iM is still present at pH 6.5, the quencher approach uncovered that the stability is not conserved at pHs higher than 6.0. This could be interpreted as a potential stabilizing effect of the iMab antibody towards the iM, allowing its folding even at pHs higher than 6.0. Interestingly, the KD obtained with the iMab approach showed a higher affinity of the antibody to the iM structrure than what published before (10). This difference could be explained by the different applied methodologies and the higher sensitivity of SwitchSENSE technology used for our assays.

Next, we observed, by several complementary biochemical techniques, that APE1 is able to stably bind the native telomeric iM structure. Differently from what observed for G4 structures, the N-terminal regions does not seem to be fundamental for the binding of APE1 to the iM structure. Indeed the obtained dissociation constants (KD) for the two proteins are in the same range (WT: 6.48 nM *vs* NΔ33: 5.46 nM). 2D NMR confirmed that the most important residues for iM binding are in the canonical duplex DNA binding region, concentrated within and around α-helices 7 and 9, in the APE1 catalytic domain. Moreover, we were able to observe real-time binding of APE1 to telomeric iM structures through Lumicks C-Trap Optical Tweezer. As we used the SMADNE approach and cell extracts obtained from a recently developed A549 cell line expressing endogenously-mEGFP-tagged APE1 protein, we could observe APE1 binding to this structure in a context similar to the physiological one, in the presence of potential competitors for both APE1 interaction and substrate binding. Moreover, contrary to the traditional SMADNE, in which cells are usually transfected with plasmids to have transient overexpression of a fluorescently labeled protein, our A549 cl.31 cell line expressed comparable levels of APE1 to A549 wild-type, excluding the potential bias due to protein overexpression. Additionally, our model was deeply characterized (see Supplementrary Fig. S3) and well represents the physiological wild-type one. Our results provide the first evidence to our knowledge that iM setup is compatible with C-Trap, as well as support the feasibility of SMADNE analysis using endogenously labelled proteins. Taken together, these findings for the first time visualize APE1 engaging with iMs.

To explore APE1 DNA repair activity of iM, we firstly examined if the AP-site containing versions of C-NAT can actually fold into a iM. Indeed, AP-sites are the primary repair target of APE1 and are one of the most frequent lesions in cells. Then, we synthetized four additional ODNs, with lesions positioned in different positions: the AP14 and AP20 ODNs bear an AP-site in the second and in the third loop, respectively, both substituting an adenine base, and AP16 and AP17 ODNs hold the AP-site located in the core of the iM structure (Fig. 3A), specifically the first and middle cytosine of the third C-run. As expected from the work of Dvoráková (32), single lesions in both loops and in the core of the iM did not disrupt the ODNs folding ability, as confirmed by CD analysis (Fig. 3B). Interestingly, both CD melting curves and the SwitchSENSE quencher approach revealed that, even at pH 5.5 all ODNs can fold into iM, the ones with the modifications in the cytosine core are less stable. Indeed, AP16 and AP17 showed significantly lower melting temperatures (AP16: 33°C, AP17: 28°C) compared to the other iM structures (C-NAT: 46°C, AP14 and AP20: 44°C) and are denatured at lower pH values (AP16: 5.8, AP17: 5.7). Again, the iMab antibody could bind all these iMs at pH 6.5, supporting a potential stabilizing effect of the antibody towards the iM, that allows their folding besides the presence of DNA modifications and at higher pHs.

Moreover, we evaluated APE1 binding ability with respect to the damaged ODNs by employing different biochemical assays. SWB and SwitchSENSE analysis revealed that both APE1 and the N-terminal truncated mutant can bind damaged iM with comparable KD compared to the C-NAT (Fig. 3F-H). Interestingly, SwitchSENSE analysis revealed a higher binding affinity of APE1^NΔ33^ towards the damaged ODNs compared to APE1^WT^. A possible interpretation for this higher affinity of the APE1 mutant towards the damaged iM structures is that, without the N-terminal unstructured region, the protein is only able to bind the iM but cannot aid the unfolding of the structure and consequently expose the AP-site for its processing, residing longer on the iM and resulting in an higher KD.

Next, we investigated APE1 cleavage activity on AP-sites present in iM telomeric sequences. We observed a strong dependence of the APE1 endonuclease activity on the AP-site position in the iM structure, as AP-sites in the core were cleaved more efficiently than the ones in the loop. AP16 and AP17 formed less stable iMs compared to AP14 and AP20, which could indicate that they can be more easily unwinded to single strand DNA by APE1, favoring the exposure of the AP-site to the processing activity by APE1. Moreover, AP16 and AP20 are in a configuration more similar to a duplex, being part of the cytosine core, and this could better resemble APE1’s canonical target. In general, the repair activity of APE1 towards AP-sites in the iM structure was around 1300 times lower compared to canonical duplex, considering the higher quantity and incubation timing of the reactions. The enzymatic activity was also from three to seven times lower compared to a single stranded unstructured substrate. Moreover, we observed that the APE1 truncated mutant was not able to cleave any of the damaged iM substrates, while it cleaved each respective duplex to a comparable extent. Interestingly, APE1’s N-terminal region seems not fundamentally important for the binding of the protein towards the iM. However, it appears to be necessary for the cleavage activity. Moreover, APE1^NΔ33^ showed a higher affinity than APE1^WT^ for iMs bearing AP-sites. APE1 might bind but not cleave AP-sites within the iM with a regulatory role independent from its canonical DNA repair function. The absence of its N-terminal domain could enhance this functional rewiring. This behaviour could be compared to APE1’s interaction with 8-oxoguanine containing G4 structures, where the damage, secondary DNA structures, and repair proteins contribute to an epitranscriptional regulatory mechanism to control gene expression.

Examination of the APE1 protein-protein interactome (42) highlights that APE1 interacts with several hnRNP proteins that are known from previous studies to influence iM folding (e.g. hnRNP K, hnRNP A1, PCBP1 (9, 24, 25)). Indeed, iM structural dynamics can be modulated by a specific protein family, called PCBPs (11, 12, 20–22). Between these proteins, we specifically focused our attention on PCBP1, as it has already been implicated in the competing formation of iM and G4 structures and in genome stability maintenance (23). We confirmed this protein-protein interaction in HeLa and U2OS cellular lines by PLA. By *in vitro* cleavage assay, we observed that PCBP1 inhibits APE1 cleavage on telomeric iM sequence, probably further stabilizing the iM structure and preventing APE1 unwinding or by strongly competing with APE1 for the substrate, as suggested by the SWB assay analyses (Fig. 5). This competition could be a potential mechanism to inhibit the repair AP-sites within iMs, which consequently could have potential unforeseen regulatory roles.

Lastly, we examined APE1-PCBP1 interplay at telomeric level in ALT-U2OS model. In the last years, the role of APE1 in telomeric regions has become clearer, as it has been observed that this protein is required for telomere maintenance in both normal and cancer cells. The depletion of APE1 has been linked with senescence and premature aging phenotypes and telomeric defects have been observed in cells using either telomerase-dependent lengthening or the ALT pathway. Indeed, APE1 telomeric localization is important for the binding of TRF2 to telomeric sequences. APE1 depletion leads to telomere uncapping and increased DDR activation at chromosome ends. By employing TRF analysis, we observed that the depletion of APE1 and PCBP1 dysregulates telomere length with opposite trends in ALT-U2OS model. ALT cells are characterized by long and variable telomeres, due to the fact that they use Homologous Recombination (HR) to elongate them. Previous studies were not able to determine telomere length differences in U2OS due to their initial extensive length. We tried to bypass this problem by sonicating the DNA before digesting it with frequently cutting restriction enzymes. The assay revealed that APE1 KO U2OS cells have slightly longer telomeres compared to the wild-type cell line, which could be related to an increased HR activation to compensate APE1 absence. Instead, PCBP1-depletion in both WT and KO models was associated with decreased telomere length. This opposite behaviour may indicate that APE1 and PCBP1 cooperate to maintain telomere length homeostasis when both proteins are present.

Previous studies suggest that APE1 and PCBP1 contribute to telomere maintenance by regulating shelterin components. An *in vitro* study published in 2012 investigated how shelterin proteins POT1, TRF1 and TRF2 impacted on BER activity at telomeres (51). Using purified proteins and DNA substrates, they observed that POT1, TRF1 and TRF2 physically interacted with APE1, FEN1 and Pol𝛽. Specifically, they observed that the shelterin complex could: i) modulate APE1 cleavage and binding activites on telomeric double stranded sequences; ii) increase FEN1 endonuclease activity on flap substrates and iii) increase Ligase I binding to help ligation. Furthermore, it was demonstrated that APE1 co-localizes with TRF2 and POT1 in cells (43), while PCBP1 interacts with TRF2 and TRF1 (14). TRF1 and TRF2 bind double-stranded telomeric DNA, with TRF2 preventing ATM-mediated DNA damage signaling (44). POT1 protects the G-rich overhang from degradation. Loss of TRF2 or POT1 from telomeres indicates dysfunction (44). Notably, APE1 depletion reduces TRF2 localization at telomeres without affecting its overall cellular levels (43). To further investigate the roles of APE1 and PCBP1 at telomeres, we examined their interaction with TRF2. Surprisingly, we observed that APE1 and PCBP1 can mutually and dependently interact with the shelterin complex protein TRF2: the absence APE1 or PCBP1 impedes the interaction of the other with TRF2. As the levels of TRF2 were not altered after APE1 or PCBP1 depletion, we can hypothesize that the proteins might form a trimeric complex or that TRF2 might alter its localization upon the depletion of these proteins. Further studies are needed to deeply characterize this aspect.

To explore APE1-iM interplay dynamics in a cellular model, we observed that the abscence of APE1 is associated with a reduction of iMab foci formation in U2OS cells. This is evidence that APE1 is involved in the regulation of iM formation *in vivo*.

APE1 and PCBP1 depletion in U2OS cells activates DDR, highlighting their role in genome stability. Genomic instability, a hallmark of cancer (52), often results from impaired DNA repair and leads to mutation accumulation (52, 53). Cancer cells bypass surveillance mechanisms like apoptosis and senescence, enabling uncontrolled growth (52, 53). To maintain telomere length and support limitless replication, they use strategies such as telomerase activation (54) or the ALT pathway (55). As introduced before, U2OS cellular model uses ALT, in which telomere elongation is independent from telomerase activity and relies on HR. APE1 depletion did not induce the phosporylation of γH2AX but induced p21 upregulation, while PCBP1 upregulated both. Thus we can speculate that they play at different levels of the ATM-p53-p21 axis in this specific model. APE1 could alter p21 expression by directly inducing p53 activation, without necessarily inducing an increased number or double-strand breaks. PCBP1 depletion instead could lead to increased DNA damage, with a upstream effect on the pathway. When both proteins are depleted, we observe reduced phosporilation of γH2AX and p21 levels, which could indicate a possible radical alteration of DNA damage signaling when they are both depleted.

In conclusion, our study not only provides a detailed characterization of an underexplored aspect of APE1, beyond the BER pathway and canonical DNA repair but, due to the high abundance of iM structures in the human genome, also opens new avenues for translational applications in biology and medicine that go beyond telomere biology.

## DATA AVAILABILITY

The data underlying this article are available in Open Science Framework (OSF), at https://doi.org/10.17605/OSF.IO/DV24S. Large kymograph data are available from the corresponding author upon request. All study data are included in the article and/or supporting information.

## SUPPLEMENATRY DATA

Supplementary Data are available at *NAR* Online.

## AUTHOR CONTRIBUTIONS

Alessia Bellina: Investigation, Methodology, Formal Analysis, Data Curation, Validation, Visualization, Writing - original draft, Writing – review & editing. Matilde Clarissa Malfatti: Conceptualization, Project administration, Supervision, Writing - original draft, Writing – review & editing. Tobias Obermann: Investigation, Methodology, Validation, Writing – review & editing. Kayla Mae Grooms: Investigation, Validation. Andreas Gjøsæther: Investigation, Validation. Zahraa Othman: Investigation, Validation. Gilmar Salgado: Investigation, Validation. Daniela Marasco: Investigation, Validation, Writing – review & editing. Antonella Virgilio: Investigation, Validation, Writing – review & editing. Veronica Esposito: Investigation, Validation. Giulia Antoniali: Methodology, Writing – review & editing. Catia Mio: Methodology. Matteo Pivetta: Methodology. Magnar Bjørås: Resources, Supervision. Barbara van Loon: Resources, Supervision, Writing – review & editing. Gianluca Tell: Conceptualization, Funding acquisition, Project administration, Supervision, Writing - original draft, Writing – review & editing.

## FUNDING

The research leading to these results has received funding from AIRC under IG 2024 - ID. 30399 project – P.I. Tell Gianluca”. The work was also supported through the support of the Departmental Strategic Plan (PSD) of the University of Udine-Interdepartmental Project on Healthy Ageing (2020-25), by additional grants from the University of Udine (‘Bando Ricerca Collaborativa’ granted by European Community - NextGenerationEU) and from the Consorzio Interuniversitario Biotecnologie - C.I.B. – (MUR-PRIN2022: “L’INNOVAZIONE DELLE BIOTECNOLOGIE NELL’ERA DELLA MEDICINA DI PRECISIONE, DEI CAMBIAMENTI CLIMATICI E DELL’ECONOMIA CIRCOLARE”) to G.T. This work was also partly supported by a grant from MUR-PRIN2022 (grant#20224F7P9Y) to G.T.. Zahraa Othman is recipient of the Excellence Eiffel fellowship by France’s Ministry of Europe and Foreign Affairs and the University of Lebanon.

## CONFLICT OF INTEREST

The authors declare no competing interests.

## Notes

https://doi.org/10.17605/OSF.IO/DV24S

### Competing Interest Statement

The authors have declared no competing interest.

