## Supplementary material for "Apurinic/apyrimidinic endodeoxyribonuclease 1 contributes to the repair of damaged intercalated-motif of telomeric sequences"

### MANUSCRIPT TITLE

**Supplementary Methods**

**RNA extraction and quantitative Reverse Transcriptase-PCR (qRT-PCR)**

RNA isolation was performed on 1.5×$10^6$ A549 wild-type cells and A549 APE1-GFP cells using the "NucleoSpin® RNA" kit (Machery-Nagel; 740955.250) according to the manufacturer's instructions. One microgram of total RNA was reverse transcribed using the SensiFAST cDNA synthesis kit (Bioline, London, UK), according to the manufacturer's instructions. The following sequences of primers were used: *APEX1* For: 5'- CCTGGACTCTCTCATCAATACTGG-3', *APEX1* Rev: 5'-AGTCAAATTCAGCCACAATCACC-3', *GAPDH* For: 5'- CCCTTCATTGACCTCAACTACATG-3', *GAPDH* Rev: 5'- TGGGATTTCCATTGATGACAAGC-3'. qRT-PCR was performed with a CFX96 Real-Time System (Bio-Rad) using SensiFAST SYBR No-ROX kit (Bioline, London, UK).

**Immunofluorescence and live-cell imaging**

For immunofluorescence 80000 cells were fixed in 4% paraformaldehyde, then washed with PBS 1× and permeabilized with 0.25% Triton X-100 in PBS 1× for 5 min. After washing with PBS 1× and blocking for 1 h with 10% FBS in Washing Buffer (10 mM Tris HCl pH7.4, 150 mM NaCl and 0.01% Tween 20), cells were incubated with APE1 primary antibodies (Novus; NB 100-101) diluted 1:100 in blocking solution for 3 h at 37 °C. After several washes in Washing Buffer 1×, cells were incubated with labelled secondary antibodies Alexa Fluor® 488 (goat anti-rabbit IgG 111-545-003 Jackson ImmunoResearch, West Grove, PA, USA) 1:400 in blocking solution for 2 h at room temperature. Cells were washed and mounted using *FluoroshieldTM with DAPI* (Sigma; 1002788770) and fixed with polish. Cells were visualized through a Leica TCS SP8 laser-scanning confocal microscope (Leica Microsystems, Wetzlar, Germany).
For Live-cell imaging, 1× $10^4$ A549 wild-type and selected clone cells were plated in an 8-well chamber (Nunc LabTek chamber slide, Merck) and grown at 37°C and 5% CO2 using *RPMI 1640* complemented medium. After 24 hours each well was washed with *PBS* and stained using *Hoechst-RPMI 1640* complemented medium and left to incubate for one hour at 37°C. Afterwards, cells were washed and visualized through a 100X objective of a Leica TCS SP8 laser-scanning confocal microscope (Leica Microsystems, Wetzlar, Germany).

**Cell cycle analysis**

Total 1× $10^6$ cells were harvested and resuspended in 1 ml of cold 70% ethanol in phosphate-buffered saline (PBS) at -20°C for 1 hour. Fixed cells were washed three times to remove ethanol and permeabilized with 100 µL PBS Triton X-100 0.1%, 10 µg/mL RNase A and 50 µg/mL Propidium Iodide in the dark for 30 minutes. Cells were then analyzed with "Attune NxT® Flow Cytometer" machine. A minimum of 3× $10^4$ cells were measured for each experimental condition. Analyses of the results were performed using FlowJo.

**Cell viability and proliferation assay**

Cell viability was measured using the 3 (4 5 dimethylthiazol 2 yl) 5 (3-carboxymethoxyphenyl) 2 (4-sulfophenyl) 2*H*-tetrazolium salt (MTS) assay (Celltiter 96 Aqueous One solution cell proliferation assay, Promega) on cells grown in 96-well plates. In detail, 4000 cells were plated on 96-wells and were allowed to attach to the plate for 24 h. The day after, cells were treated separately with different amounts of *MMS* (Merck KGaA; 129925) for 8 hours, or *CDDP* (Merck KGaA, P4394), for 24 hours. After treatment, the MTS solution was added to each well, and the plates were incubated for 2 h at 37 °C. Absorbance was measured at 490 nm using a multiwell plate reader (Synergy H1, Agilent). All experiments were run at least in triplicate. The values were standardized to wells containing media alone and the cell viability was expressed as a fold change compared to the DMF-treated cells.

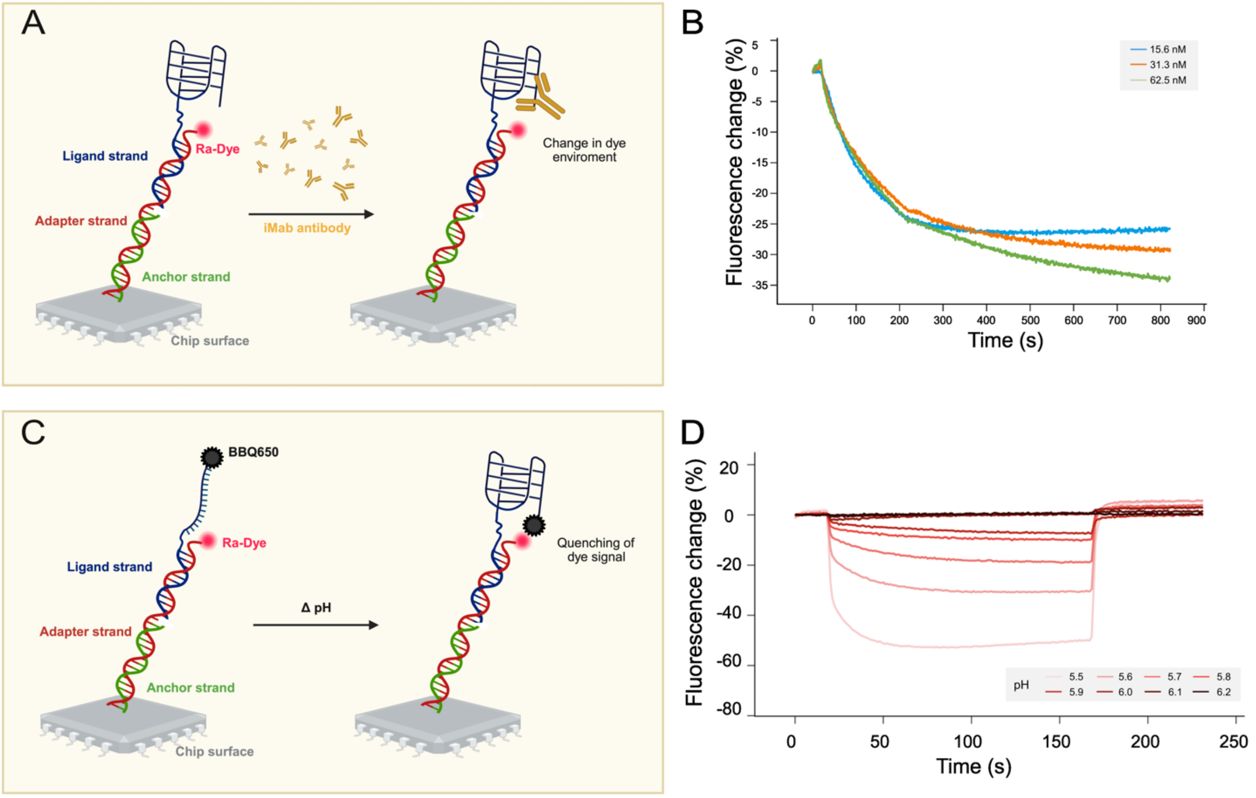


**Figure Supplementary 1.** A) Schematic representation of i-motif recognition by iMab antibody performed on SwitchSense. The anchor strand is indicated in green, the adapter strand in red, derivatized at its end with Ra-Dye, and the ligand strand in blue, with the iM structure at its end. The binding of the iMab antibody induces a change in the dye environment. B) Real-time fluorescence signals and fits of a representative experiment measuring the association and dissociation phases of iMab at different concentrations towards immobilized C-NAT at pH 5.5. Time (expressed in s) and Fluorescence change (expressed in %) are reported on the x- and y- axis, respectively. C) Schematic representation of i-motif formation assay through the quencher approach, performed on SwitchSense. The anchor strand is indicated in green, the adapter strand in red, derivatized at its end with Ra-Dye, and the ligand strand in blue, with the iM structure at its end, derivatized with BBQ650. The iM folding is monitored through the injection of buffers with increasing pH and by measuring the dye fluorescence. D) Real-time fluorescence signals of C-NAT derivatized with a quencher, after the injection of buffers with increasing pHs. Time (expressed in s) and Fluorescence change (expressed in %) are reported on the x- and y- axis, respectively. Curves referring to buffers with different pHs are represented in different colors.

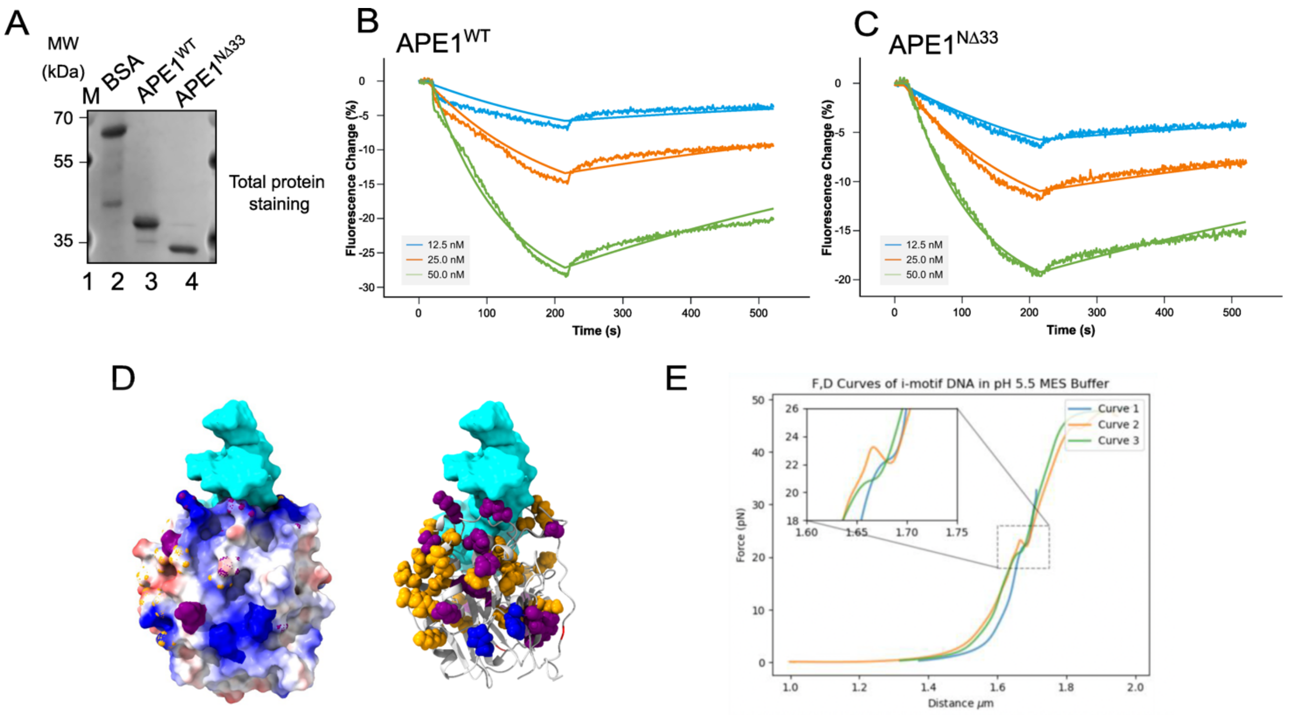


**Figure Supplementary 2.** A) Total protein staining showing the loading of equal amounts of the recombinant proteins BSA, APE1$^{WT}$, APE1$^{N\Delta 33}$. On the sides, the electrophoretic marker (M) is loaded, and the different molecular weights (MW) are indicated and expressed in kDa. B-C) Real-time fluorescence signals and fits of a representative experiments measuring the association and dissociation phases of APE1$^{WT}$ (B) and APE1$^{N\Delta 33}$ (C) at different concentrations towards immobilized C-NAT at pH 5.5. Time (expressed in s) and Fluorescence change (expressed in %) are reported on the x- and y- axis, respectively. D) Depiction of the side view of NMR-derived binding analysis on APE1 structure (PDB code: 1bix), relative to Figure 2D. E) Force-distance curves (n=3) of the 3x-C-NAT substrate. The unfolding events are zoomed in. Distance (expressed in μm) and Force (expressed in pN) are reported on the x- and y- axis, respectively.

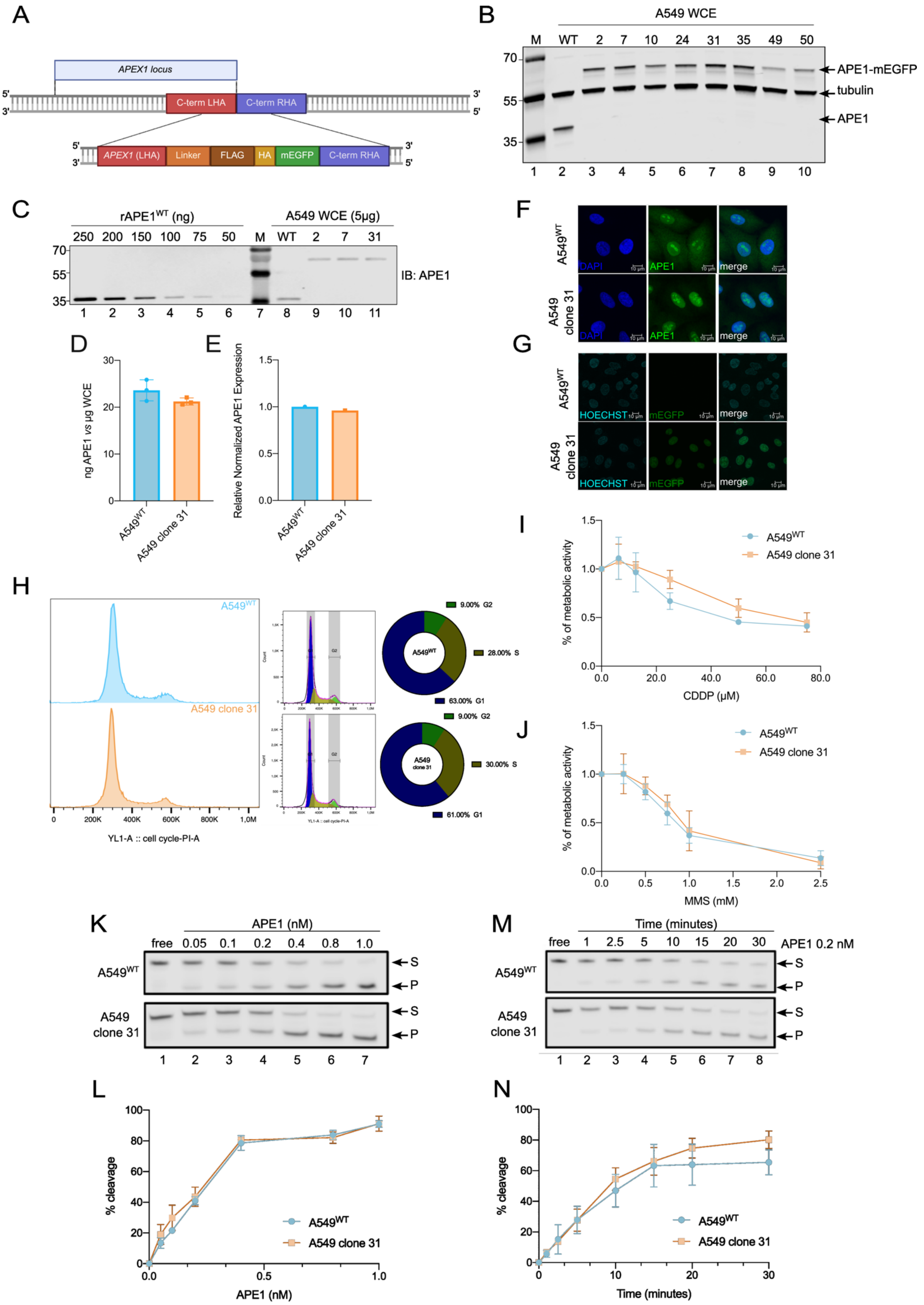
A
APEX1 locus
5'
3'
C-term LHA
C-term RHA
APEX1 (LHA)
Linker
FLAG
HA
mEGFP
B
A549 WCE
M WT 2 7 10 24 31 35 49 50
70
55
35
APE1-mEGFP
tubulin
APE1
1 2 3 4 5 6 7 8 9 10
C
rAPE1WT (ng)
250 200 150 100 75 50
M
A549 WCE (5μg)
WT 2 7 31
IB: APE1
1 2 3 4 5 6 7 8 9 10 11
D
ng APE1 vs μg WCE
A549WT
A549 clone 31
E
Relative Normalized APE1 Expression
F
A549WT
A549 clone 31
DAPI
APE1
merge
G
HOECHST
mEGFP
H
A549WT
A549 clone 31
YL1-A :: cell cycle-PI-A
9.00% G2
28.00% S
63.00% G1
30.00% S
61.00% G1
I
% of metabolic activity
CDDP (μM)
J
MMS (mM)
K
APE1 (nM)
free 0.05 0.1 0.2 0.4 0.8 1.0
S
P
1 2 3 4 5 6 7
M
Time (minutes)
free 1 2.5 5 10 15 20 30
APE1 0.2 nM
1 2 3 4 5 6 7 8
L
% cleavage
APE1 (nM)
N
Time (minutes)

**Figure Supplementary 3.** A) Schematic representation of CRISPR–Cas9 technology used to generate the endogenous C-term tagging of APE1 in A549 cells. B) Representative western blot analysis comparing APE1-mEGFP levels to APE1$^{WT}$. 20 µg of A549 WCE were loaded. Tubulin was used as a loading control. C-D) Quantification of APE1-mEGFP *via* rAPE1$^{WT}$ standard curve. Protein quantification was estimated on a biological triplicate using 5 µg of A549 WCE and serial rAPE1$^{WT}$ dilutions (250-50 ng). E) APE1-mEGFP gene expression quantification. *APEX1* expression levels were quantified *via* qPCR on a technical replicate and normalized to *GAPDH*. F-G) Immunofluorescence and live-cell imaging analysis for evaluating APE1-mEGFP localization. Both A549$^{WT}$ and Clone 31 cells were fixed and stained with α-APE1 488 antibody and DAPI (F). For live-cell imaging, cells were stained with Hoechst (G). The µm scale is reported on the bottom left. H) Cell sorting analysis for Clone 31 cell-cycle phases quantification. G1, S and G2 phases distribution was measured on Clone 31 via Propidium Iodide (PI) using A549$^{WT}$ as control. Percentage distributions are shown in the donut charts. I-J) Measurement of Clone 31 viability via MTS assay. Increasing concentrations of Cisplatin (CDDP) (I) and Methyl Methane Sulfonate (MMS) (J) were administered for both clone 31 and A549$^{WT}$ to evaluate metabolic activity. DMF and RPMI 1640 were used as control vehicles, respectively. K-L) Representative AP-site incision assay and related estimation of APE1-mEGFP endonuclease activity. Increasing concentrations of APE1 (0.05-1.0 nM) from WCE of A549$^{WT}$ and Clone 31 were tested to calculate endonuclease activities on a biological triplicate. The percentage of cleavage activity was estimated by comparing the Product (P) signal to the sum of both Substrate (S) and P signals. M-N) Representative AP-site incision assay and related estimation of APE1-mEGFP endonuclease kinetics. 0.2 nM APE1 from WCE of A549$^{WT}$ and Clone 31 was incubated with substrate for increasing time points (1-30 minutes) to estimate endonuclease kinetics on a biological triplicate. The percentage of cleavage activity was estimated by comparing the Product (P) signal to the sum of both Substrate (S) and P signals.

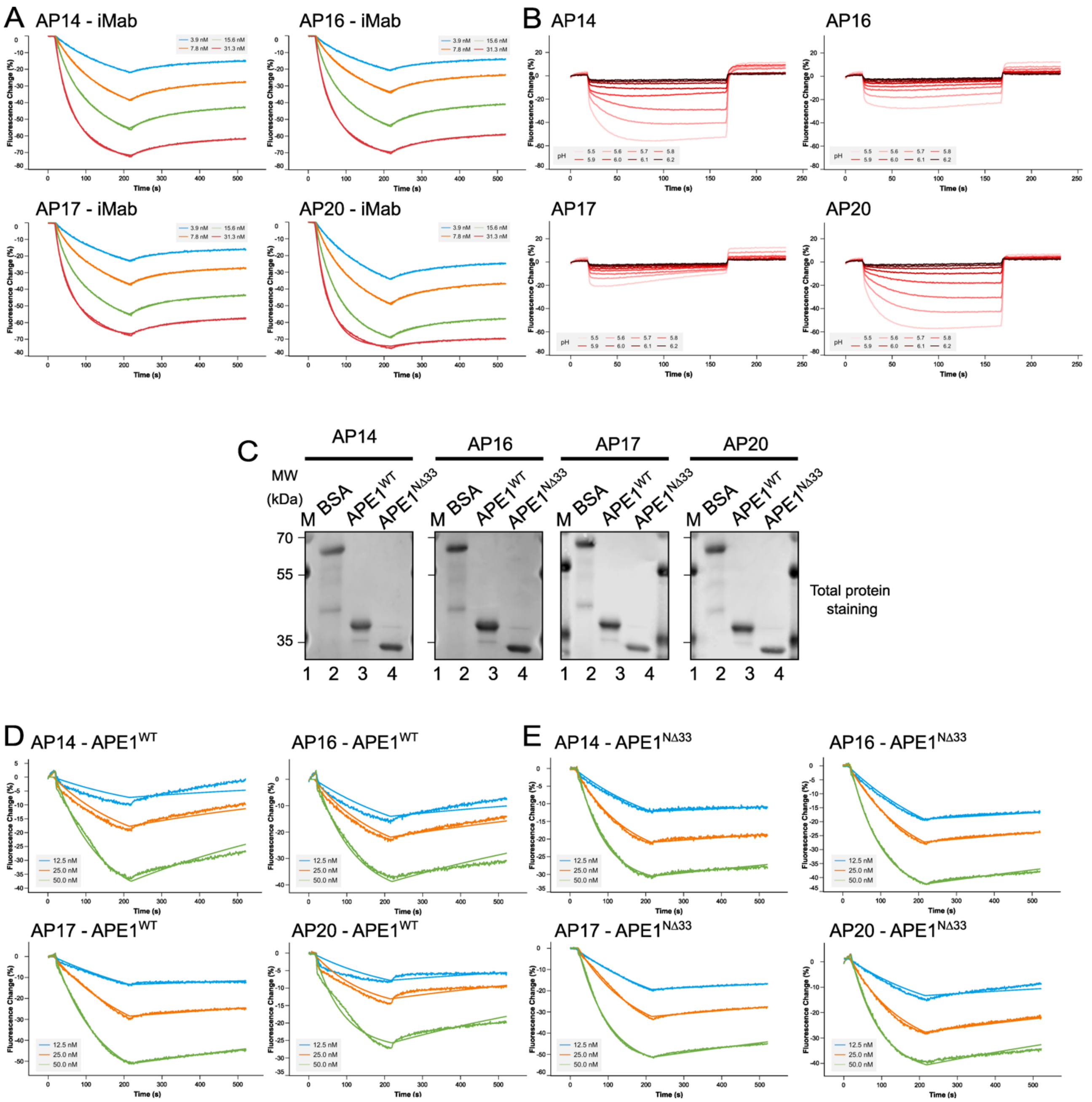


**Figure Supplementary 4.** A) Real-time fluorescence signals and fits of a representative experiment measuring the association and dissociation phases of iMab at different concentrations towards immobilized AP14, AP16, AP17 and AP20 at pH 6.5. Time (expressed in s) and Fluorescence change (expressed in %) are reported on the x- and y- axis, respectively. B) Real-time fluorescence signals of AP14, AP16, AP17 and AP20 derivatized with a quencher, after the injection of buffers with increasing pHs. Time (expressed in s) and Fluorescence change (expressed in %) are reported on the x- and y- axis, respectively. Curves referring to buffers with different pHs are represented in different colors. C) Total protein staining showing the loading of equal amounts of the recombinant proteins BSA, $APE1^{WT}$, $APE1^{N\Delta33}$. On the sides, the electrophoretic marker (M) is loaded, and the different molecular weights (MW) are indicated and expressed in kDa. D-E) Real-time fluorescence signals and fits of a representative experiment measuring the association and dissociation phases of $APE1^{WT}$ (D) and $APE1^{N\Delta33}$ (E) at different concentrations towards immobilized AP14, AP16, AP17 and AP20 at pH 6.5. Time (expressed in s) and Fluorescence change (expressed in %) are reported on the x- and y- axis, respectively.

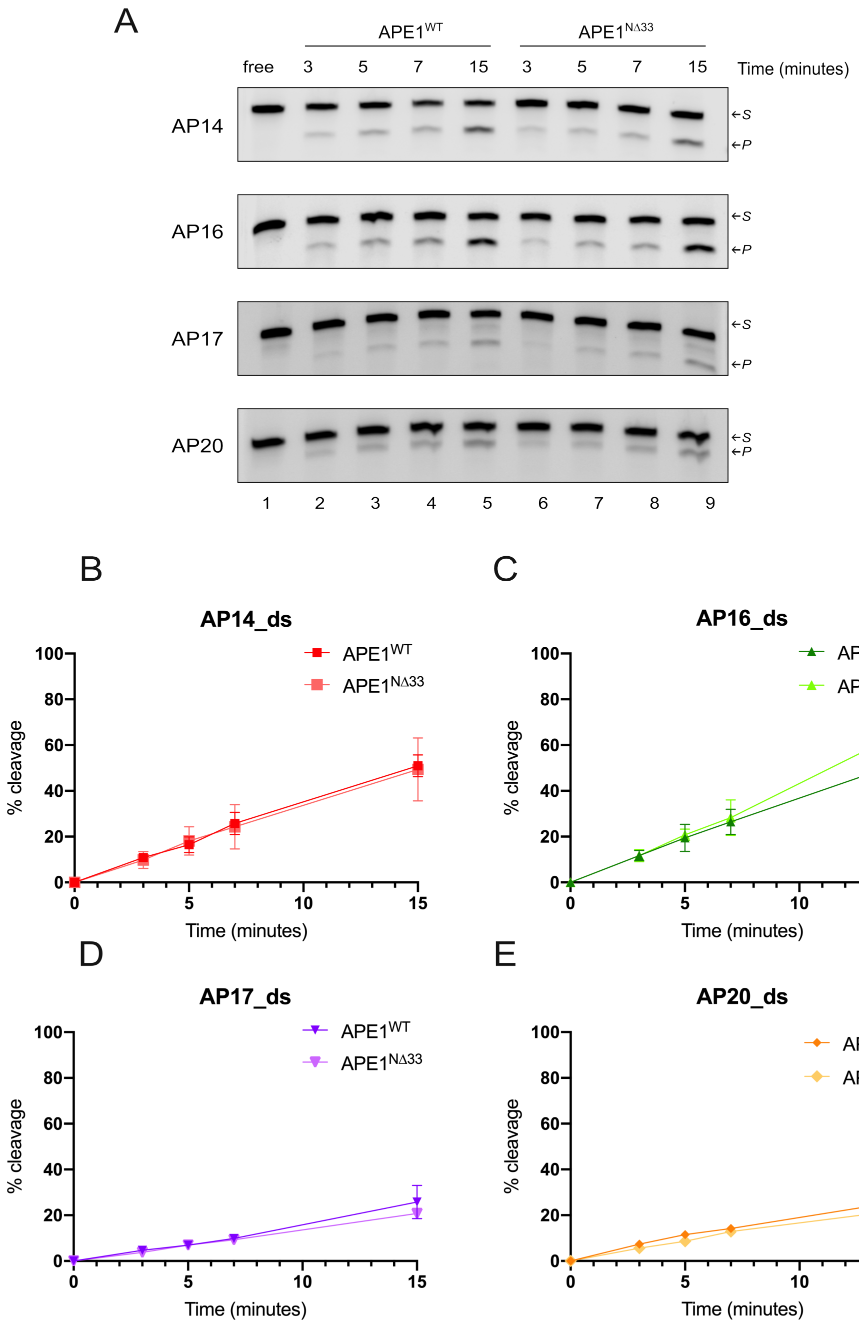


**Figure Supplementary 5.** A) Representative denaturing polyacrylamide gels of cleavage analysis obtained on all ds_ substrates alone (lane 1, "free"), with APE1$^{WT}$ protein (lanes 2-5) and with APE1$^{N\Delta33}$ protein (lanes 6-9). On the right, the substrate and the product bands are indicated by two arrows. A constant dose of APE1$^{WT}$ or APE1$^{N\Delta33}$ (0.125 nM) was incubated with the each oligonucleotide at 37°C, and the reactions were stopped at different time points, indicated upon the gel and expressed in minutes. B-E) Relative graph illustrating the time-course kinetics activity of APE1$^{WT}$ and APE1$^{N\Delta33}$ recombinant proteins on AP14_ds (B), AP16_ds (C), AP17_ds (D) and AP20-ds (E). Time (minutes) and percentage of cleavage (%) are reported on the x- and y- axis, respectively. Data are expressed as mean ± SD of three independent technical replicas.

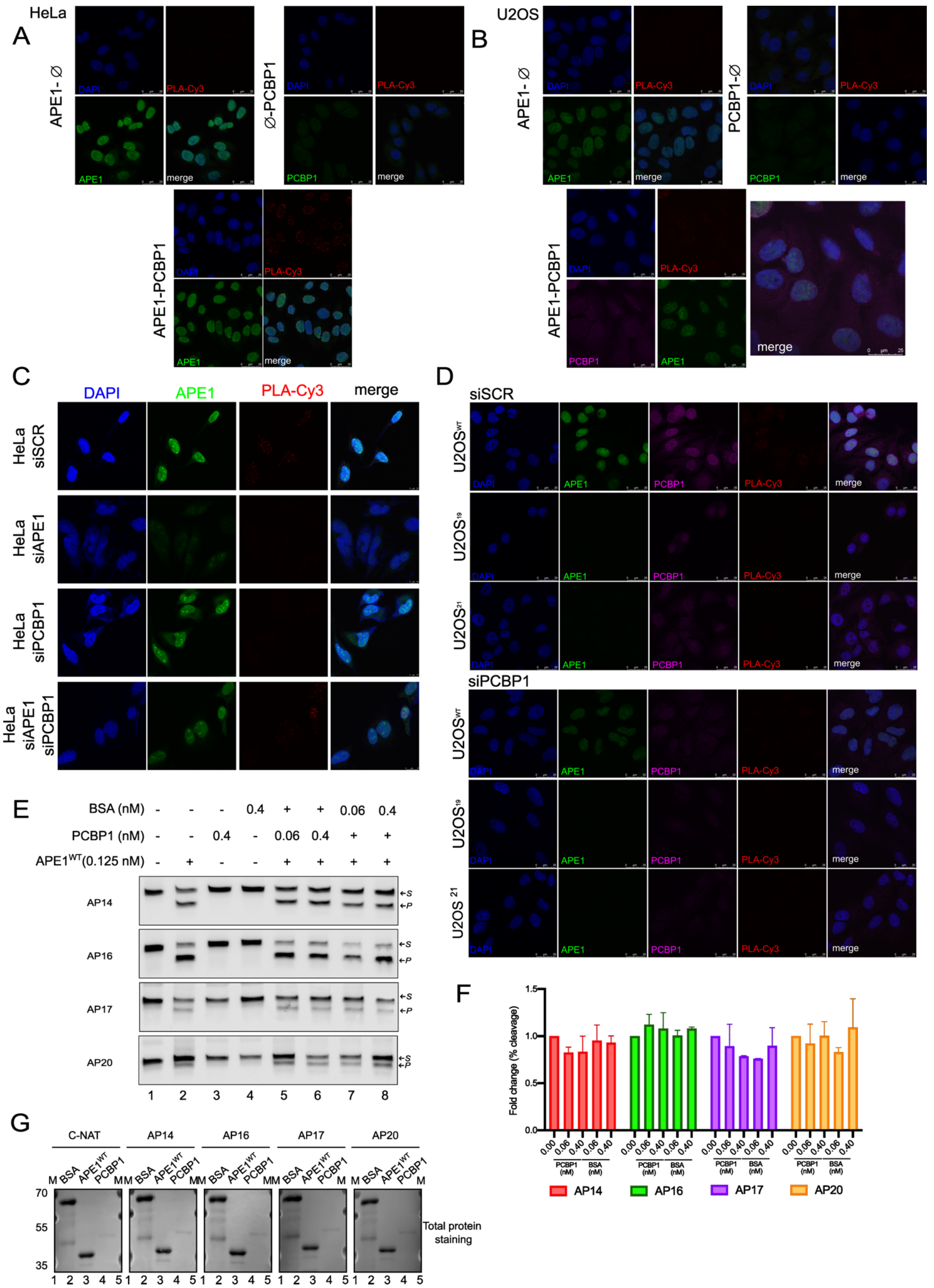


**Figure Supplementary 6.** A) PLA analysis between APE1 and PCBP1 proteins in HeLa cells (APE1-PCBP1). Technical negative controls of PLA were obtained by omission of the antibody directed *versus*

PCBP1 (APE1-Ø) or *versus* APE1 (Ø-PCBP1). APE1 staining was obtained with rabbit-488 (green) secondary antibody, while the nuclear staining, obtained with DAPI, is in blue. PLA dots are in red (Cy3-555). The merge panel shows the overlay between the four channels and reports the scale bar expressed in µm. B) PLA analysis between APE1 and PCBP1 proteins in U2OS cells. Technical negative controls of PLA were obtained by omission of the antibody *versus* PCBP1 (APE1-Ø) or *versus* APE1 (Ø-PCBP1). APE1 and PCBP1 staining were obtained with rabbit-488 (green) and mouse-633 (magenta) secondary antibodies, while the nuclear staining, obtained with DAPI, is in blue. PLA dots are in red (Cy3-555). The merge panel shows the overlay between the four channels and reports the scale bar expressed in µm. C) PLA reactions for HeLa cells transiently silenced for APE1 (siAPE1), PCBP1 (siPCBP1), or both proteins. Scramble control is also reported (siSCR). PLA spots (in red) show the interaction between APE1 and PCBP1 proteins. D) PLA reactions for U2OS clones, respectively U2OS$^{WT}$, U2OS$^{19}$ and U2OS$^{21}$, transiently silenced for PCBP1. Scramble controls are also reported. PLA spots (in red) show the interaction between APE1 and PCBP1 proteins. E) Representative denaturing polyacrylamide gels of cleavage analysis obtained on all substrates alone (lanes 1), with APE1$^{WT}$ protein only (lanes 2), with PCBP1 and BSA only (lanes 3 and 4), with PCBP1 co-incubated with APE1 (lanes 5-6) and BSA co-incubated with APE1 (lanes 7-8). On the right, the substrate and the product bands are indicated by two arrows. The ODNs were pre-incubated with a variable dose of PCBP1 or BSA (upon the gel, nM) for 90 minutes at 4°C. Next, a constant dose of APE1 (0.125 nM) was added to the reactions and incubated at 37°C for 15 minutes, when the reaction was stopped. F) Relative graph illustrating the fold change of APE1$^{WT}$ recombinant protein activity after pre-incubation with different amount of PCBP1 or BSA on AP14 (red), AP16 (green), AP17 (purple) and AP20 (orange). PCBP1 and BSA concentrations (nM) and fold change of the percentage of cleavage (%) are reported on the x- and y- axis, respectively. Data are expressed as mean ± SD of three independent technical replicas. G) Total protein staining showing the loading of the recombinant proteins BSA, APE1$^{WT}$, PCBP1. On the sides, the electrophoretic marker is loaded, and the different molecular weights are expressed in kDa.

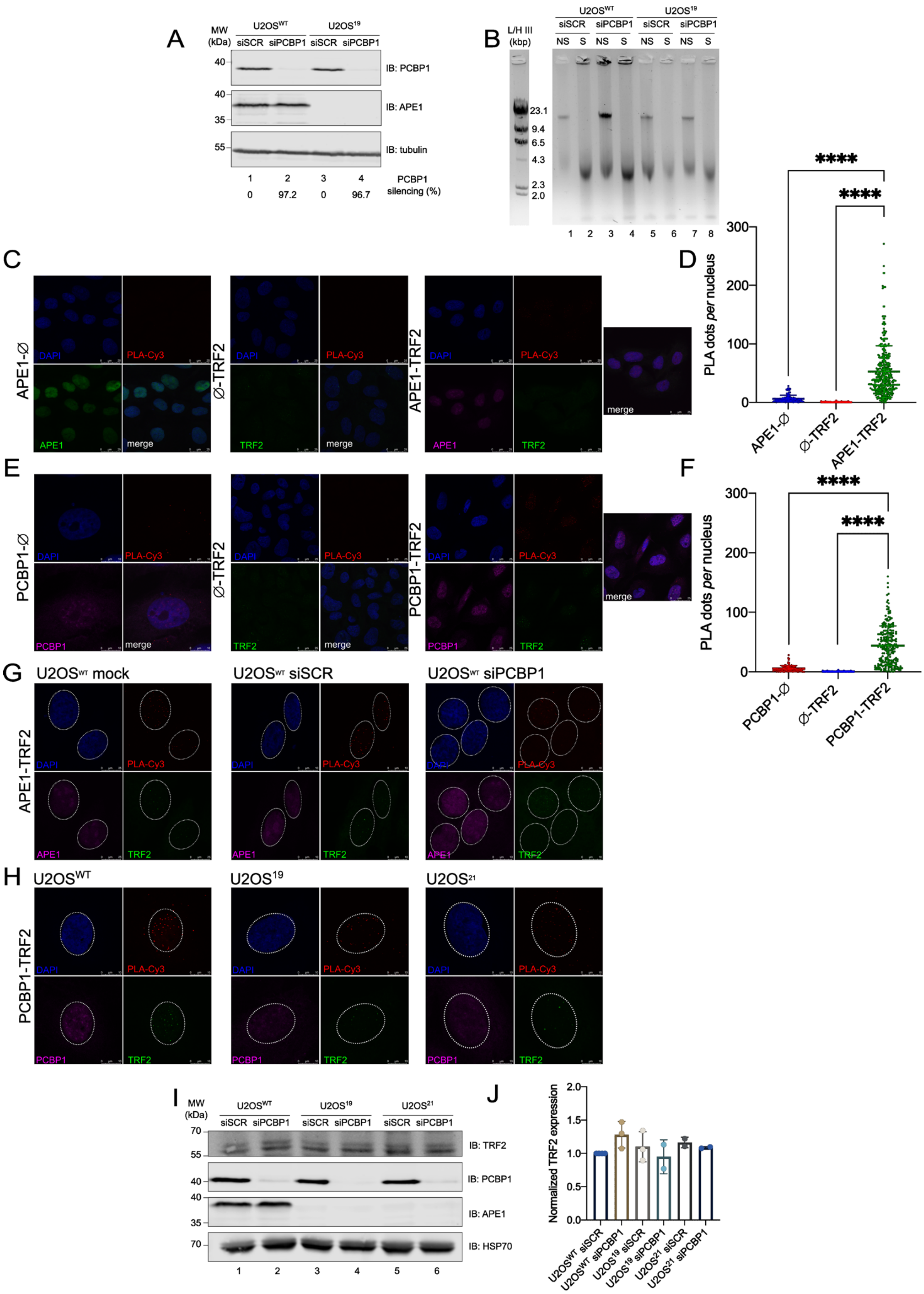

A
MW (kDa)
U2OSWT
U2OS19
siSCR
siPCBP1
IB: PCBP1
IB: APE1
IB: tubulin
PCBP1 silencing (%)
97.2
96.7
B
L/H III (kbp)
NS
S
23.1
9.4
6.5
4.3
2.3
2.0
C
APE1-Ø
Ø-TRF2
APE1-TRF2
DAPI
PLA-Cy3
APE1
TRF2
merge
D
PLA dots per nucleus
****
E
PCBP1-Ø
PCBP1-TRF2
PCBP1
F
G
U2OSWT mock
U2OSWT siSCR
U2OSWT siPCBP1
H
U2OSWT
U2OS19
U2OS21
I
IB: TRF2
IB: HSP70
J
Normalized TRF2 expression
U2OSWT siSCR
U2OSWT siPCBP1
U2OS19 siSCR
U2OS19 siPCBP1
U2OS21 siSCR
U2OS21 siPCBP1

**Figure Supplementary 7.** A) Western blot analysis of PCBP1 and APE1 levels in $U2OS^{WT}$, $U2OS^{19}$ cell lines after PCBP1 transient silencing. Tubulin was used as normalizer. Molecular weight (MW, expressed in kDa) are reported on the left. The percentage of PCBP1 silencing is reported under each lane. B) Representative agarose gel of DNA extracted from $U2OS^{WT}$, $U2OS^{19}$ cell lines after PCBP1 transient silencing, non-sonicated (NS) or sonicated (S) for four cycles. On the left, the length marker Lambda/Hind III is reported and the length of each band is indicated in kbp. C) PLA analysis between APE1 and TRF2 proteins in U2OS cells. Technical negative controls of PLA were obtained by omission of the antibody directed *versus* TRF2 (APE1-Ø) or *versus* APE1 (Ø-TRF2). TRF2 and APE1 staining were obtained with rabbit-488 (green) and mouse-633 (magenta) secondary antibodies, while the nuclear staining, obtained with DAPI, is in blue. PLA dots are in red (Cy3-555). The merge panel shows the overlay between the four channels and reports the scale bar expressed in µm. D) Graph depicting the number of TRF2-APE1 PLA dots *per* nucleus in $U2OS^{WT}$, compared to technical negative controls (APE1 only and TRF2 only). Average and standard deviation values are plotted ($n = 1$ for controls, $n = 3$ for APE1-TRF2). E) PLA analysis between PCBP1 and TRF2 proteins in U2OS cells. Technical negative controls of PLA were obtained by omission of the antibody directed *versus* TRF2 (PCBP1-Ø) or *versus* PCBP1 (Ø-TRF2). TRF2 and PCBP1 staining were obtained with rabbit-488 (green) and mouse-633 (magenta) secondary antibodies, while the nuclear staining, obtained with DAPI, is in blue. PLA dots are in red (Cy3-555). The merge panel shows the overlay between the four channels and reports the scale bar expressed in µm. F) Graph depicting the number of TRF2-PCBP1 PLA dots *per* nucleus in $U2OS^{WT}$, compared to technical negative controls (PCBP1 only and TRF2 only). Average and standard deviation values are plotted ($n = 1$ for controls, $n = 3$ for PCBP1-TRF2). Statistical analysis was performed using one-way ANOVA test. G) Single panels relative to the merge reported in Figure 6D. PLA dots are in red (Cy3-555), TRF2 and APE1 staining were obtained with mouse-488 (green) and rabbit-633 (magenta) secondary antibodies, while the nuclear staining, obtained with DAPI, is in blue. The scale is indicated and expressed in µm. H) Single panels relative to the merge reported in Figure 6F. PLA dots are in red (Cy3-555), TRF2 and PCBP1 staining were obtained with mouse-488 (green) and rabbit-633 (magenta) secondary antibodies, while the nuclear staining, obtained with DAPI, is in blue. The scale is indicated and expressed in µm. I) Western blot analysis of TRF2, PCBP1 and APE1 levels in $U2OS^{WT}$, $U2OS^{19}$ and $U2OS^{21}$ cell lines after PCBP1 transient silencing. HSP70 was used as loading control and normalizer. Molecular weight (expressed in kDa) is reported on the left. J) Densitometric analysis of TRF2 expression levels, normalized to HSP70. Fold change values relative to $U2OS^{WT}$ siSCR, arbitrary set to 1, are shown. Values are mean ± SD of two independent replicates.
